# Decoding Mechanisms of PTEN Missense Mutations in Cancer and Autism Spectrum Disorder using Interpretable Machine Learning Approaches

**DOI:** 10.1101/2025.01.16.633473

**Authors:** Miao Yang, Jingran Wang, Ziyun Zhou, Wentian Li, Gennady Verkhivker, Fei Xiao, Guang Hu

## Abstract

Missense mutations in oncogenic proteins that are concurrently associated with neurodevelopmental disorders have garnered significant attention. Phosphatase and tensin homolog (PTEN) serves as a paradigmatic model for mapping its mutational landscape and identifying genotypic predictors of distinct phenotypic outcomes, including cancer and autism spectrum disorder (ASD). Despite extensive research into the genotype-phenotype correlations of PTEN mutations, the mechanisms underlying the dual association of specific PTEN mutations with both cancer and ASD (PTEN-cancer/ASD mutations) remain elusive. This study introduces an integrative approach that combines machine learning (ML) with structural dynamics to elucidate the molecular effects of PTEN-cancer/ASD mutations. Analysis of biophysical and network biology-based signatures reveals a complex energetic and functional landscape. Subsequently, an ML model and corresponding integrated score were developed to classify and predict PTEN-cancer/ASD mutations, underscoring the significance of protein dynamics in predicting cellular phenotypes. Further molecular dynamics simulations demonstrated that PTEN-cancer/ASD mutations induce dynamic alterations characterized by open conformational changes restricted to the P loop and coupled with inter-domain allosteric regulation. This research aims to enhance the genotypic and phenotypic understanding of PTEN-cancer/ASD mutations through an interpretable ML model integrated with structural dynamics analysis. By identifying shared mechanisms between cancer and ASD, the findings pave the way for the development of novel therapeutic strategies.

## 1. INTRODUCTION

Epidemiological studies have demonstrated that certain cancers occur more frequently in individuals with neurodevelopmental disorders (NDDs), suggesting a significant correlation between these conditions^1^. Cancer originates from the dysregulation of cellular growth, proliferation, and differentiation^2^, whereas NDDs arise from disruptions in nervous system development, affecting cognitive, motor, social, and behavioral functions^3^. Despite their distinct clinical manifestations, emerging evidence has uncovered shared cellular pathways, proteins and genetic mutations underlying both conditions^4–6^. For instance, both diseases involve dysfunctions in key biological processes, including chromatin remodeling and signaling through the PI3K/mTOR and MAPK pathways^7^. Notably, over one-third of genes causally linked to cancer have also been implicated in the onset of NDDs^8^, with phosphatase and tensin homolog (PTEN) protein exemplifying this intersection^9^.

PTEN comprises several functional domains, including N-terminal PIP2-binding domain (PBD), catalytic phosphatase domain (PD), membrane-binding C2 domain (C2D), and C-terminal tail (CTT), each essential for its diverse roles in cellular signaling^10^. Missense mutations in the PTEN gene are associated with a broad spectrum of diseases^11^, encompassing various cancers and Autism Spectrum Disorder (ASD)^12^. These mutations can be categorized into three types based on clinical outcomes: (1) PTEN-caner mutations, present in patients with cancer only; (2) PTEN-ASD mutations, identified in patients with ASD only; and (3) PTEN-cancer/ASD mutations, observed in patients with both cancer and ASD^12–14^. Compared to the first two categories, PTEN-cancer/ASD mutations warrant significant attention due to their dual phenotypic associations. For example, the PTEN-cancer/ASD mutation R173C has been identified in conditions such as PTEN hamartoma tumor syndrome, Cowden syndrome, ASD, and various cancers, involving colorectal cancer and brain neoplasms, disrupting the catalytic site regulation and altering its phospho-regulation^14^and increasing domain flexibility and fluctuation in the CBR3 loop, interdomain regions, and C-terminal tail^15^, which resulted in the loss of phosphatase function and enhanced cellular transformation^16^. However, the dual structural and functional impacts of PTEN-cancer/ASD mutations remain underexplored.

Various methodologies have been employed to analyze PTEN mutations and their clinical relevance. Experimental approaches, including biophysical analyses^17^, structural and statistical assessments^18^, deep mutational scanning^19^, and phenotypic studies in model organisms^20^, have been used to investigate the functional impacts of these mutations. Nevertheless, these methods often lack scalability and fail to elucidate detailed molecular mechanisms, thereby limiting the comprehensive understanding of PTEN mutations. In contrast, computational approaches, such as molecular dynamics (MD) simulations^21^, elastic network models (ENMs)^22^, and protein structure network (PSN)-based analyses^21^ have uncovered conformational changes, long-range allosteric effects, and disruptions in PTEN function and stability. These studies have been pivotal in distinguishing PTEN-cancer from PTEN-ASD mutations, where PTEN-ASD mutations primarily induce localized dynamic increases, particularly at the active site, leading to local instability that may affect substrate binding and catalytic activity, thereby contributing to ASD pathology. Conversely, PTEN-cancer mutations result in significant global structural destabilization, including reduced active site stability and increased interdomain dynamics^23^, potentially compromising PTEN’s tumor-suppressive functions and promoting oncogenesis. Furthermore, a machine learning (ML) model^24^ has been developed to classify PTEN mutations by analyzing their molecular effects on protein structure and function. Although effective in distinguishing PTEN-cancer and PTEN-ASD mutations, the mechanistic basis of PTEN-cancer/ASD mutations and their role in the genotype-phenotype relationship remains poorly understood, necessitating further investigation.

Biophysical insights into missense mutations are critical for evaluating protein functions^25–29^. This study aims to bridge the gap between PTEN mutations and their disease phenotypes by leveraging PTEN structural dynamics, including three steps (Figure 1). First, three types of mutations were curated from gene mutation databases, and molecular and network signatures were assigned to each mutation type through multi-level analyses. Next, an ML model incorporating an integrated scoring function bases on the above features was developed to distinguish phenotypic similarities, enabling the classification of PTEN-cancer/ASD mutations. Lastly, the molecular mechanisms underlying PTEN-cancer/ASD mutations were investigated by exploring the interpretable natures of the ML model and quantifying their conformational dynamics and long-range perturbation dynamics. Taken together, the methodological advances presented here elucidate molecular mechanisms underlying pathogenic mutations involving co-occurring diseases, and thus assist in assigning novel therapeutic strategies.

**Figure 1.**
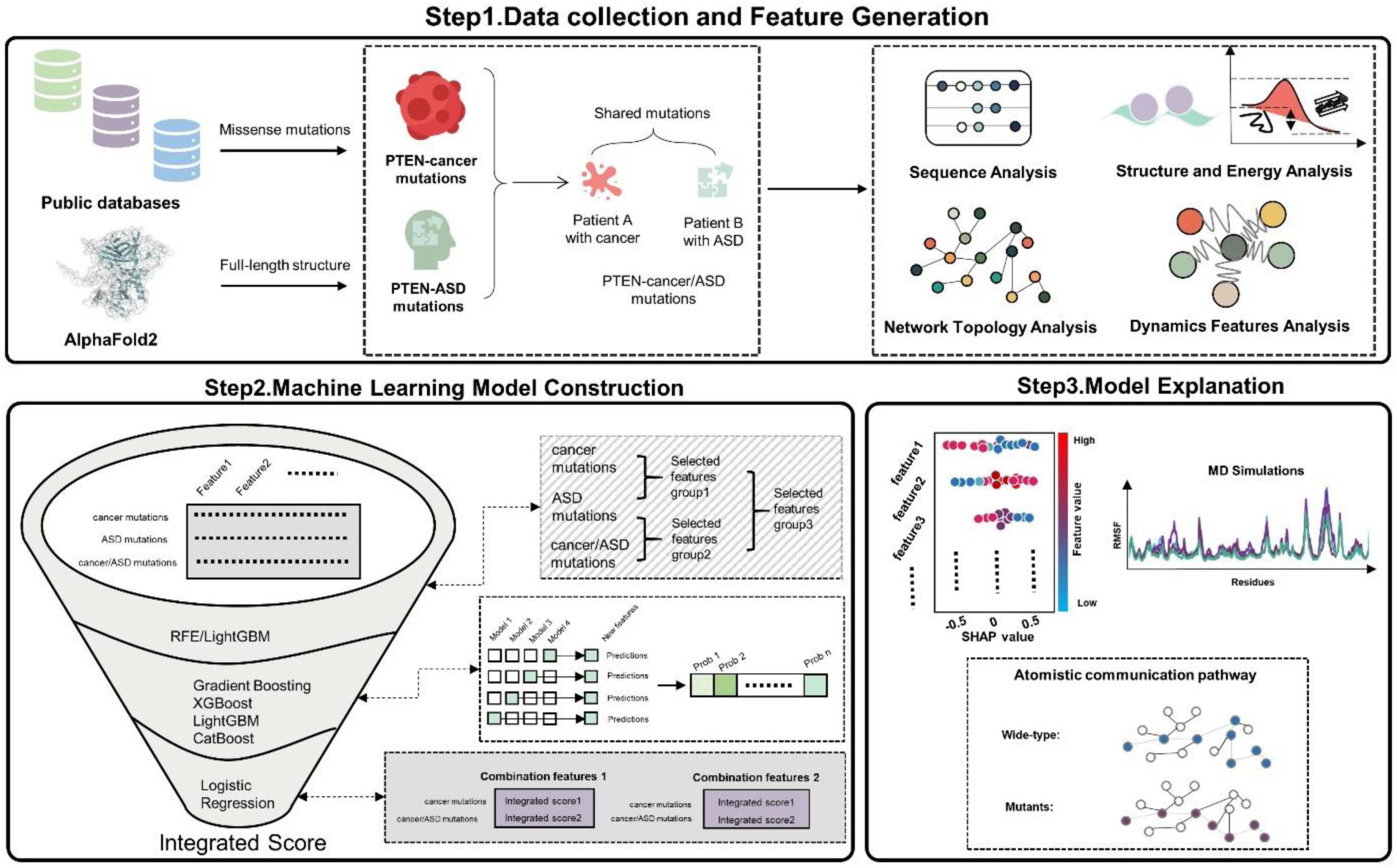
Overview of the integrative workflow for PTEN mutation classification and analysis. It consists of data collection and feature generation, machine learning model construction and model explanation. Model interpretability is achieved via SHAP-values to highlight influential features, while MD simulations and communication pathway analysis provide deeper insights into local and long-range structural effects distinguishing wild-type (WT) from mutants.

## 2. MATERIALS AND METHODS

Detailed information of structural modeling, molecular feature calculations, and MD simulations are provided in the Text S1.

### Data collection and data preparation

Three types of PTEN missense mutations including PTEN-cancer, PTEN-ASD, and PTEN-cancer/ASD mutations, as well as their phenotypic information, were systematically retrieved from the ClinVar database^30^, the Genome Aggregation database^31^, and the Cosmic database^32^using the search parameters “Missense”, “Pathogenic” and “SNVs”. To further ensure comprehensive data collection, additional PTEN mutations with corresponding phenotypic annotations from previously published studies were incorporated ^14, 17^ (Table S1).

### Molecular features calculation

A total of molecular features at sequence, structure and dynamic levels were calculated to classify PTEN missense mutations, encompassing relative Accessible Surface Area (*RASA*), conservation score, Shannon information entropy *S(i)*, mutual information (*MI*), folding Gibbs free energy (*ΔΔG*), Degree Centrality (*DC*), Clustering Coefficient (*C*), Betweenness Centrality (*BC*), Closeness Centrality (*CC*), Eigenvector Centrality (*EC*), Mean-Square Fluctuations (*MSF*), stiffness, Dynamic Flexibility Index (*DFI*), sensitivity, effectiveness.

### Machine learning framework

To construct the predictive model (Figure 2), multiple interpretable ML algorithms were employed. The framework commenced with Recursive Feature Elimination (RFE)^33^ in conjunction with the LightGBM classifier^34^ to filter features, thereby identifying those most critical for disease classification with mean precision obtained via 5-fold cross validation. RFE systematically reduces the number of features under consideration by evaluating each feature’s impact on model performance to select the optimal subset. Utilizing the feature importance rankings from LightGBM, the most effective feature combinations were identified and subsequently input into various classification models to construct a stacking model. The classification models included Gradient Boosting^35^, XGBoost^36^, LightGBM^34^, and CatBoost^37^. This approach enhanced model’ robustness and accuracy in classifying the effects of PTEN missense mutations. Stratified 10-fold cross-validation was employed to evaluate the classification performance of the models. The predicted probabilities from these models were consolidated into a probability matrix and subsequently input into a logistic regression model for final integration.

**Figure 2.**
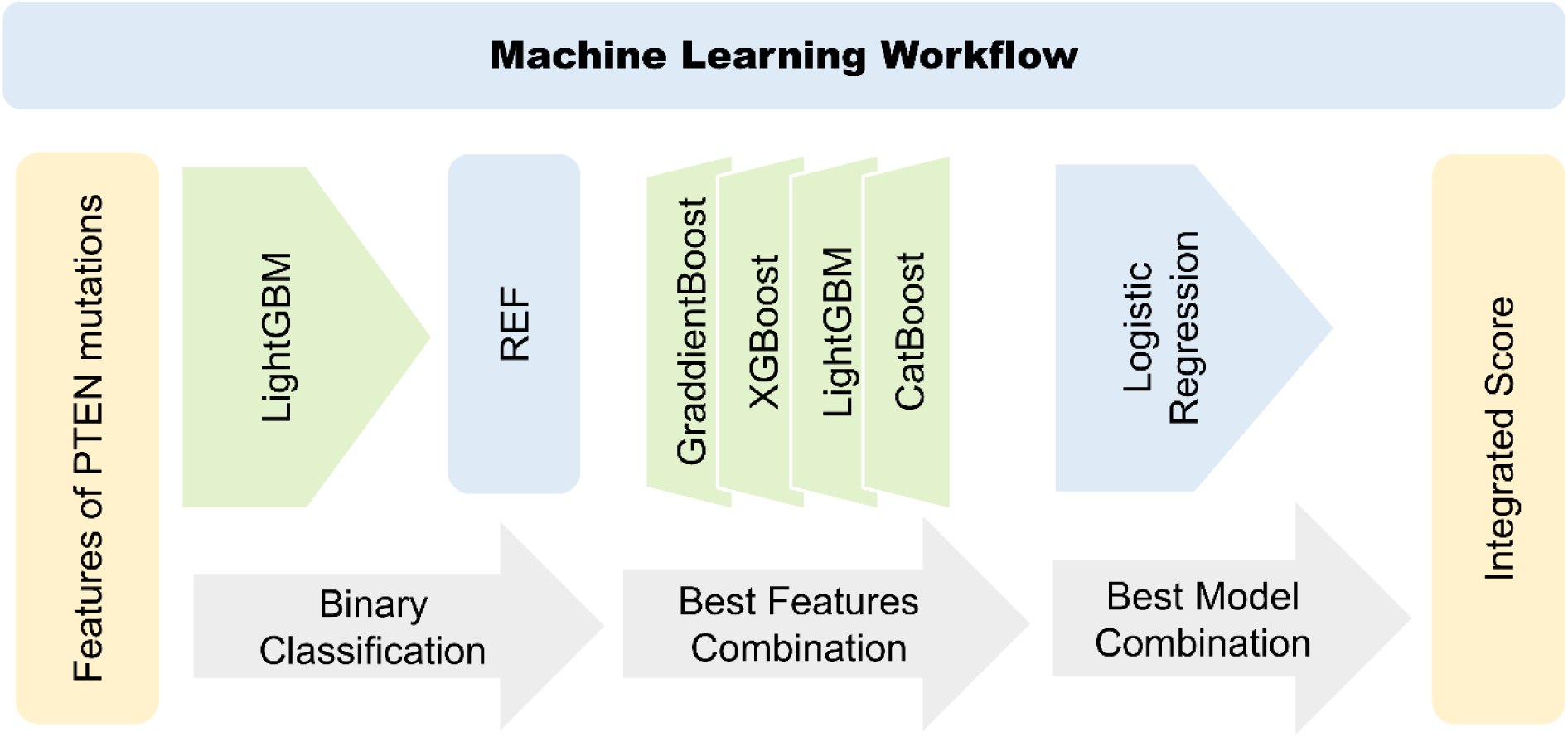
Overview of multi-feature integration and ML workflow. PTEN mutations are first extracted and then binary classified by LightGBM. The feature set is then optimised using RFE. The optimal combination of features is determined using ensemble models (GradientBoost, XGBoost, LightGBM and CatBoost). The optimal combination of models is then determined using logistic regression, and an Integrated Score is generated for each mutation with different phenotypes.

Additionally, our model develops an IS to distinguish mutations causing similar phenotypes. The score is derived from the probabilities predicted by the model and facilitates an initial assessment of the potential phenotypes associated with relevant missense mutations. For binary classification distinguishing different phenotypes, the IS ranges from 0 to 1, with mutations with IS close to 0 or 1 being more likely to correspond to their associated phenotypes. This scoring system provides an initial evaluation of unknown missense mutations, aiding their differentiation based on phenotypic impact.

### Performance analysis and model interpretability

The model’s performance was evaluated using Receiver Operating Characteristic (ROC) curves and the corresponding Area Under the Curve (AUC) values. The AUC, ranging from 0 to 1, quantifies the model’s ability to distinguish between positive and negative samples, with higher values indicating superior performance across all classification thresholds. The AUC was instrumental in determining the optimal model and parameter combinations within the stacking model. For model interpretability, Tree SHapley Additive exPlanations (SHAP)^38^ were employed to enhance both global and individual interpretability within the LightGBM framework. SHAP, grounded in cooperative game theory and inspired by Shapley values, provides a robust methodology for attributing the contribution of individual features to model predictions. On a global scale, SHAP quantifies feature importance, elucidating their aggregate impact on model behavior. At the individual level, SHAP reveals the contribution of specific features to each prediction, offering a granular view of the decision-making process. By leveraging SHAP, comprehensive insights into the interplay of features were attained, enriching both the global and individual interpretability of the model.

### Communication pathway analysis

Based on MD simulations (see Text S1), two methodologies were employed to explore the allosteric influences of mutations. Dynamic residue network (DRN) analysis was performed using the MD-TASK software^39^. In this approach, network nodes were defined by the *C*α and *C*β atoms extracted from the MD trajectories, and edges were established when the interatomic distance ≤ 6.5 Å. The optimal allosteric pathways were identified by evaluating the product of edge occurrence frequencies along the path and site betweenness derived from the set of shortest paths. Additionally, the Floyd-Warshall algorithm was utilized to delineate the shortest edges connecting mutational sites to allosteric sites, thereby revealing the allosteric communication pathways^40^.

## 3. RESULTS AND DISSCUSSION

### PTEN-cancer/ASD mutations are preferentially buried, located in the Phosphatase Domain, and exhibit higher conservation

A total of 541 missense mutations were collected and categorized based on their phenotypic associations into three groups: PTEN-cancer (409 mutations), PTEN-ASD (79 mutations), and PTEN-cancer/ASD (53 mutations) (Figure 3a, Figure S1). These mutations were mapped onto the PTEN protein structure (Figure 3b, Figure S1), revealing distinct localization patterns across the groups. Specifically, PTEN-cancer mutations were predominantly enriched in the ATP-binding Type A region, situated between the P loop and the WPD loop. In contrast, PTEN-ASD mutations were distributed across the ATP-binding Type A and surrounding regions, with a subset extending into the ATP-binding Type B region. Notably, the PTEN-cancer, PTEN-ASD, and PTEN-cancer/ASD mutations predominantly cluster within the PD, whereas relatively fewer mutations occur in the PBD, C2D, and CTT (Figures 3c-e). Further analysis of the *RASA* (Figures 3f-h and Table S2) indicated differential solvent accessibility among the mutation groups. Specifically, 63% of PTEN-cancer mutations, 72% of PTEN-ASD mutations, and 81% of PTEN-cancer/ASD mutations were situated in buried or partially exposed regions. This trend underscores a higher propensity for PTEN-cancer/ASD mutations to occur in less solvent-accessible areas of the protein.

**Figure 3.**
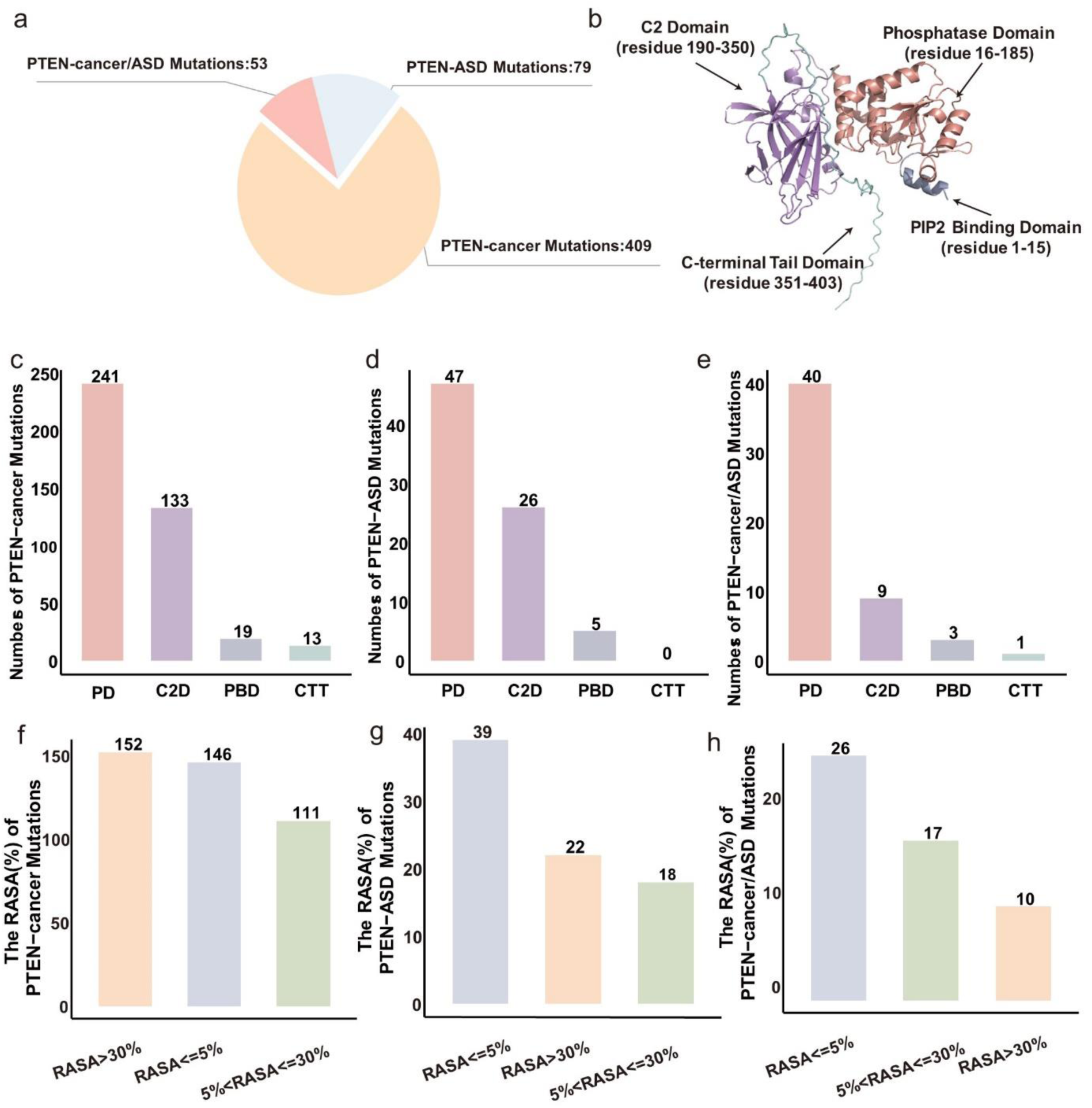
Landscape of PTEN mutation data. (a) Statistical distribution of PTEN missense mutations categorized into three phenotypic groups: PTEN-cancer (orange), PTEN-ASD (blue), and PTEN-cancer/ASD (red). (b) Full-length AlphaFold structure of PTEN highlighting its functional domains: PBD, PD, C2D, and CTT domain. (c) PTEN-cancer, (d) PTEN-ASD, (e) PTEN-cancer/ASD groups’ distribution across the PBD, PD, C2D, and CTT. (f) PTEN-cancer, (g) PTEN-ASD, (h) PTEN-cancer/ASD groups’ *RASA* calculation results.

To elucidate the sequence characteristics underlying the three types of PTEN mutations, three sequence-based signature including conservation score, *S(i)*, and *MI* were calculated. Using the conservation score threshold of ≥ 5 to define conserved residues, 215 out of 403 residues (53.34%) in the PTEN sequence were identified as conserved (Figure S2). As depicted in Figure 4a, conserved residues were mutated in 68% of the PTEN-cancer group (183/242), 81% of the PTEN-ASD group (60/67), and 83% of the PTEN-cancer/ASD group (37/43) (Table S3). This distribution indicated a higher propensity for PTEN-cancer/ASD mutations to occur at conserved sites. According to *S(i)*, 63% of conserved residues (259/409) were mutated in the PTEN-cancer group, 73% (58/79) in the PTEN-ASD group, and 81% (43/53) in the PTEN-cancer/ASD group (Figures 4b, Table S3). Co-evolutionary analysis revealed that PTEN-cancer/ASD mutations exhibit lower co-evolutionary scores compared to PTEN-cancer and PTEN-ASD mutations (Figure 4c, Table S3). This suggests that PTEN-cancer/ASD mutations may have a more significant impact on protein function due to their occurrence in residues that are less evolutionarily constrained. Statistical comparisons of conservation score, *S(i)*, and *MI* among the mutation groups revealed significant differences between the PTEN-cancer and PTEN-ASD groups, as well as between the PTEN-cancer and PTEN-cancer/ASD groups (Figures 4d-f).

**Figure 4.**
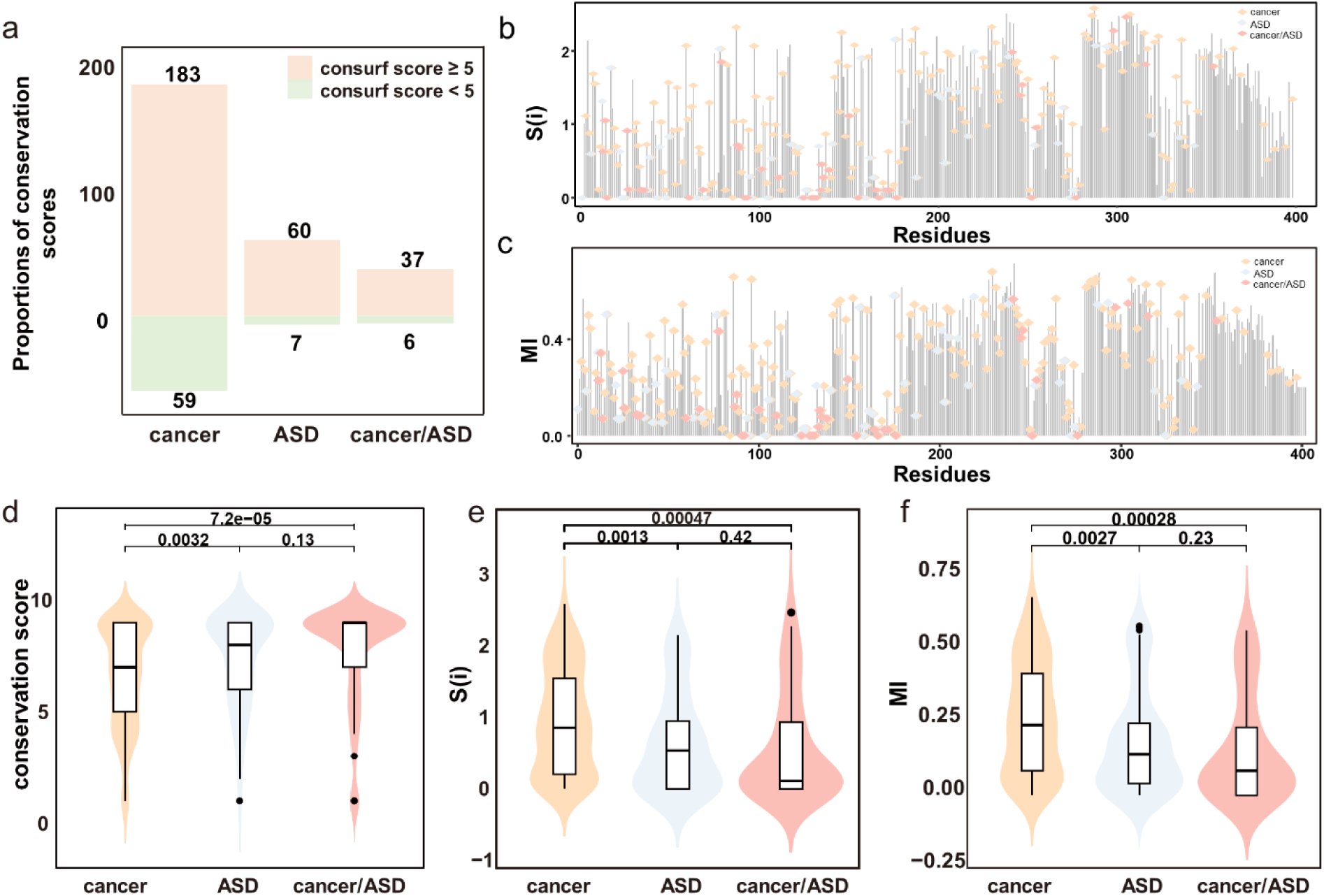
Sequence landscape of PTEN mutations. (a) Proportions of conservation scores for PTEN missense mutations. (b) *S(i)*, and (c) *MI* values for three types of PTEN mutations. (d-f) Significant differences (P-values<0.05) were observed in all three sequence features between the PTEN-cancer group and the other two groups.

### Distinct Energetic Landscapes of PTEN-cancer/ASD mutations

To elucidate the thermodynamic determinants of mutation hotspots and their functional relevance, FoldX^41^ was employed to construct mutants for each mutation group and calculate the local free energy changes induced by these mutations (Table S4). As depicted in Figures 5a and 5b, compared with the mutations related to ASD, significant energy changes could be observed between the PTEN-cancer/ASD and the cancer mutations. Specifically, the most pronounced energy changes were observed in mutations localized to the PD and the C2D (Figure 5c),which were particularly enriched in critical functional regions, such as the P loop and TI loop^13^. The top 10 PTEN-cancer/ASD mutations with the largest increase in free energy include C136R (*ΔΔG*=12.456 kcal/mol), I135R(*ΔΔG*=12.252 kcal/mol), G132D(*ΔΔG*=11.151 kcal/mol), T26P(*ΔΔG*=9.521 kcal/mol), G165V(*ΔΔG*=8.964 kcal/mol), F241S(*ΔΔG*=6.012 kcal/mol), D252G(*ΔΔG*=5.086 kcal/mol), M35V(*ΔΔG*=4.846 kcal/mol), I135K(*ΔΔG*=4.720 kcal/mol), Y68N(*ΔΔG*=4.5985 kcal/mol), which may induce severe disturbances in intramolecular dynamics or impair the interactions^23, 42, 43^ Among them, I135R and C136R, located in the PD, are classified as severe pathogenic mutations implicated in both cancer and ASD^23^. Similarly, the G165V mutation, situated in the TI loop, is identified as a loss-of-function variant, further highlighting its detrimental effect on PTEN function^43^. These findings underscore the substantial thermodynamic and functional impact of PTEN-cancer/ASD mutations, particularly those affecting key structural regions essential for PTEN activity and regulation.

**Figure 5.**
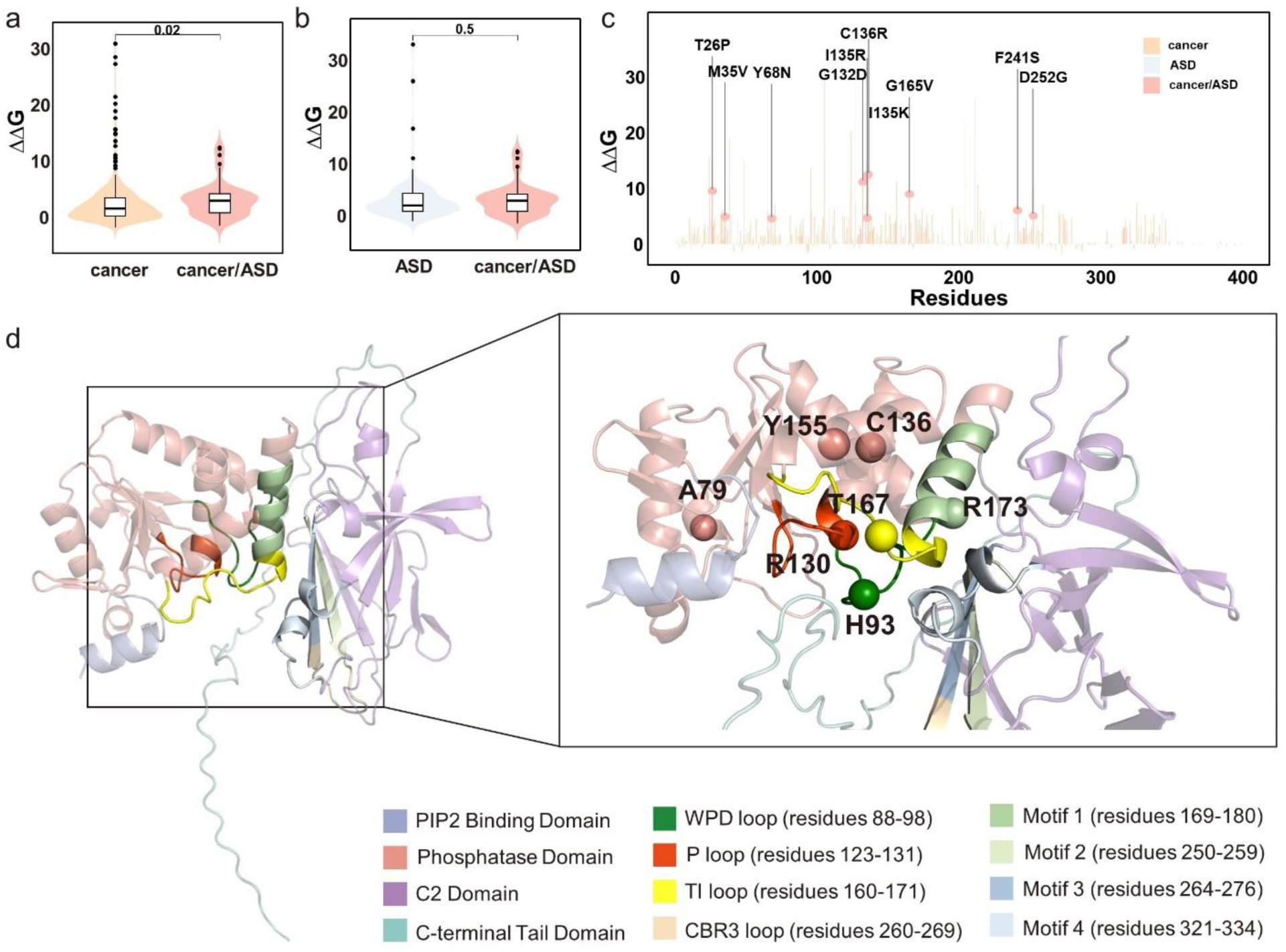
Energetic landscapes of PTEN-cancer/ASD mutations. Significance test analysis of free energy change between PTEN-cancer/ASD (red) and (a) PTEN-cancer mutations (yellow) as well as (b) PTEN-ASD mutations (blue) respectively. (c) Distribution of free energy changes in the PTEN protein sequence for PTEN-cancer, PTEN-ASD mutations, and PTEN-cancer/ASD mutations. The top 10 PTEN-cancer/ASD mutations with the highest free energy changes are marked in red font. (d) Detailed visualization of specific mutations and their corresponding free energy changes in the PTEN sequence.

However, based on our energy analysis, other key PTEN-cancer/ASD mutations, such as R130Q(*ΔΔG*=1.827 kcal/mol), R173C(*ΔΔG*=3.115 kcal/mol), Y155C(*ΔΔG*=4.054 kcal/mol) did not exhibit significant energy increases with only C136R(*ΔΔG*=12.456 kcal/mol) showing a substantial energy increase as shown in Figure 5c and 5d. R130Q, and Y155C are located within the ATP-binding type A region of the active site, whereas the R173C is located at the interdomain interface of Motif 1^23^. Additionally, some mutations were found to enhance rather than destabilize protein stability. For instance, the H93R mutation (*ΔΔG*=-1.430 kcal/mol), located in WPD loop, is strongly associated with both cancer and ASD, and its importance has been highlighted in previous studies^42, 44^. Similarly, the Y167N mutation (*ΔΔG*=-0.009 kcal/mol), located near Motif 1, exhibited increased protein stability in our energy analysis, and has been reported to induce long-range conformational changes, significantly perturbing intramolecular dynamics^23^. Moreover, the A79T mutation (*ΔΔG*=0.132 kcal/mol) located near the P loop, has been linked to potential functional disruption of PTEN^45^.

Our free energy profiles support previous findings that strong hotspot mutations lead to a cancer phenotype, while weak or moderate mutations are associated with NDDs^46^. However, challenges remain in revealing the complex landscape of energy changes induced by PTEN-cancer/ASD mutations, as their heterogeneous consequences may depend on the cellular context and dynamics.

### The network and dynamic landscape of PTEN-cancer/ASD mutations

To systematically assess the structural and dynamics impacts of PTEN mutations, we employed an amino acid contact energy network and computed five topological features for each mutation. The differences in network centrality metrics between mutant and WT proteins were calculated to characterize the topological changes induced by PTEN mutations (Figure 6, Table S5).

**Figure 6.**
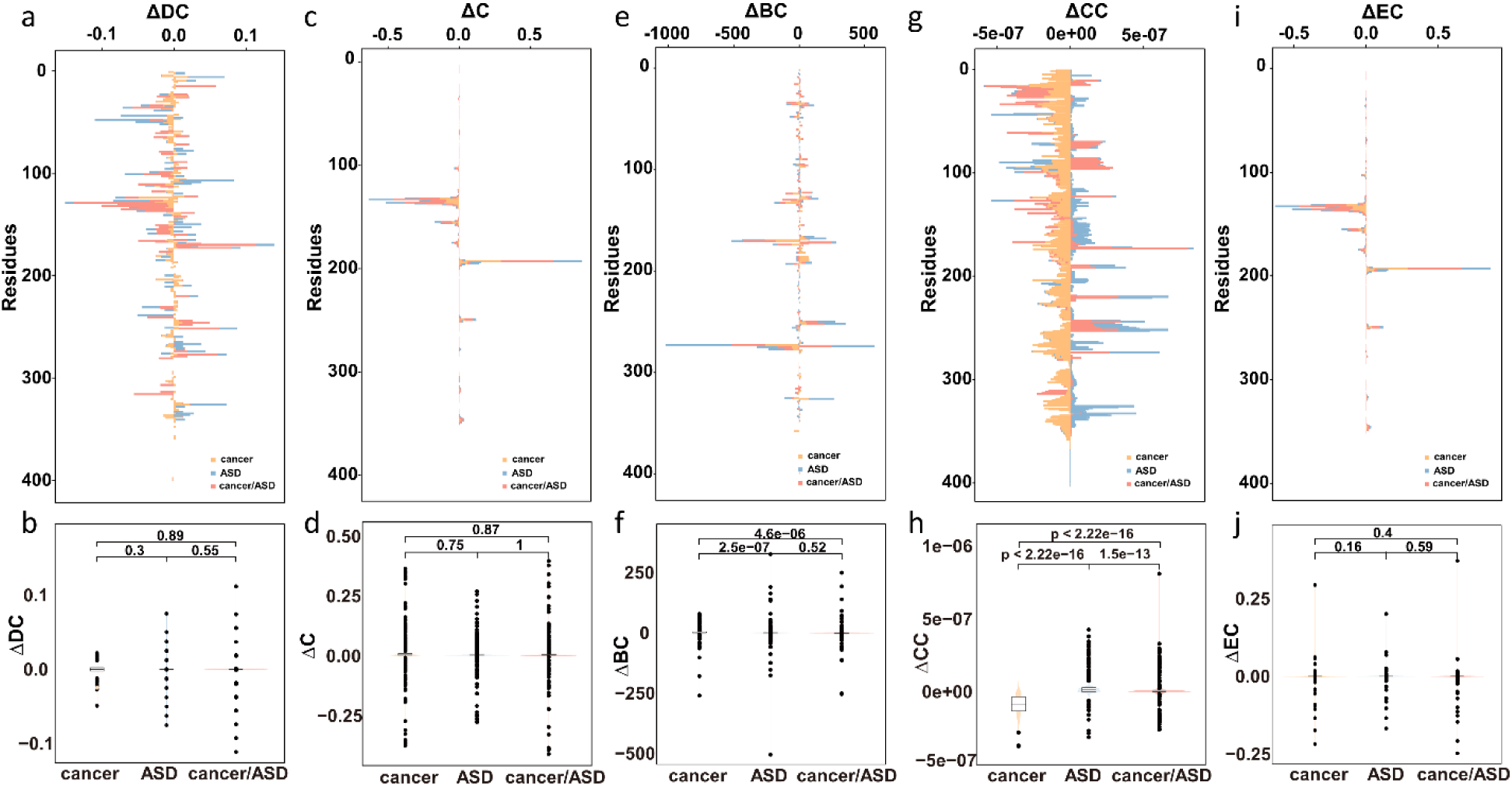
Network analysis of PTEN mutations. Distribution of the difference in (a) Δ*DC*, (c) Δ*C*, (e) Δ*BC*, (g) Δ*CC*, (i) Δ*EC*, respectively, between the mutated and the WT. Statistical analysis of (b) Δ*DC*, (d) Δ*C*, (f) Δ*BC*, (h) Δ*CC*, (j) Δ*EC*, respectively, across groups.

Examination of Δ*DC* revealed varied topological alterations among the groups. The PTEN-cancer/ASD cohort exhibited a broader distribution of Δ*DC* compared to the PTEN-cancer and PTEN-ASD groups, indicating differential degrees of network centrality perturbation (Figure 6a). However, no significant differences in Δ*DC* were observed across the groups (Figure 6b). Similarly, the assessment of the Δ*C* demonstrated no statistically significant variations among the PTEN-cancer, PTEN-ASD, and PTEN-cancer/ASD groups (Figures 6c and 6d), which suggests that clustering behavior within the network remains relatively unaffected across the different PTEN mutation groups. In contrast to Δ*DC* and Δ*C*, Δ*BC* analysis highlighted distinct network disruptions (Figures 6e and 6f). The PTEN-cancer/ASD group displayed a dispersed Δ*BC* distribution, indicating intermediate topological changes compared to the pronounced alterations in critical residues observed in the PTEN-cancer group and the relatively stable profile with fewer outliers in the PTEN-ASD group. Moreover, statistical network analysis revealed significant differences in Δ*BC* between the PTEN-cancer and PTEN-ASD groups (P-value=2.5e^-7^) as well as between the PTEN-cancer and PTEN-cancer/ASD groups (P-value=4.6e^-6^). Notably, the PTEN-cancer/ASD group exhibited the highest Δ*BC* among all cohorts, whereas no significant difference was observed between the PTEN-ASD and PTEN-cancer/ASD groups. Furthermore, the evaluation of Δ*CC* indicated substantial dispersion within the PTEN-cancer/ASD group compared to the PTEN-cancer and PTEN-ASD groups (Figures 6g and 6h). Statistical comparisons confirmed significant differences in Δ*CC* among all group pairings: PTEN-cancer vs. PTEN-ASD (P-value<2.22e^-16^), PTEN-cancer vs. PTEN-cancer/ASD (P-valuee<2.22e^-16^), and PTEN-ASD vs. PTEN-cancer/ASD (P-value=1.5e^-13^). Specifically, the PTEN-cancer/ASD mutations exhibited a reduction in Δ*CC*, highlighting notable alterations in network closeness centrality within this group. In contrast, the analysis of Δ*EC* did not reveal significant differences among the three groups (Figures 6i and 6j), suggesting that the influence of individual residues within the overall network remains consistent across the different PTEN mutation contexts.

Given the significant distinctions observed in Δ*BC* and Δ*CC* between the PTEN-cancer/ASD group and the PTEN-cancer or PTEN-ASD groups, further analysis was conducted to elucidate the underlying structural implications. Residues exhibiting elevated Δ*BC* changes in the PTEN-cancer/ASD group were primarily located at interdomain interfaces and in the vicinity of the P loop. In contrast, mutations within the PTEN-cancer group were predominantly observed around the TI loop and at the junction between the PD and C2D, while those in the PTEN-ASD group were concentrated at interdomain interfaces. Additionally, the mutations associated with the most substantial decreases in Δ*CC* (A34V, I135R, I135K, H61R, H61Y) were all localized within the PD. Notably, H61R and H61Y are situated in the ATP Binding Type A motif, whereas I135R and I135K are located in the WPD loop, a region critical for substrate catalysis. These findings underscore the differential impact of PTEN mutations on network centrality metrics and further corroborate our previous results regarding the structural and functional significance of PTEN-cancer/ASD mutations^23^.

We performed ENM modeling to investigate the dynamic effects of PTEN-cancer/ASD mutations (Table S6). Distinct trends were observed in the effectiveness, *DFI*, stiffness, *MSF*, and sensitivity metrics for the three mutation types, particularly in their distribution within critical functional regions of the PD (Figure 7a-7e). Our analysis revealed that PTEN-cancer/ASD mutations are characterized by higher effectiveness, *DFI*, and stiffness values, coupled with lower *MSF* and sensitivity values. These mutations predominantly cluster within the ATP Binding Motif Type A, especially around the P loop, within the PD. In contrast, PTEN-ASD and PTEN-cancer mutations exhibit broader distributions, including regions such as the WPD loop.

**Figure 7.**
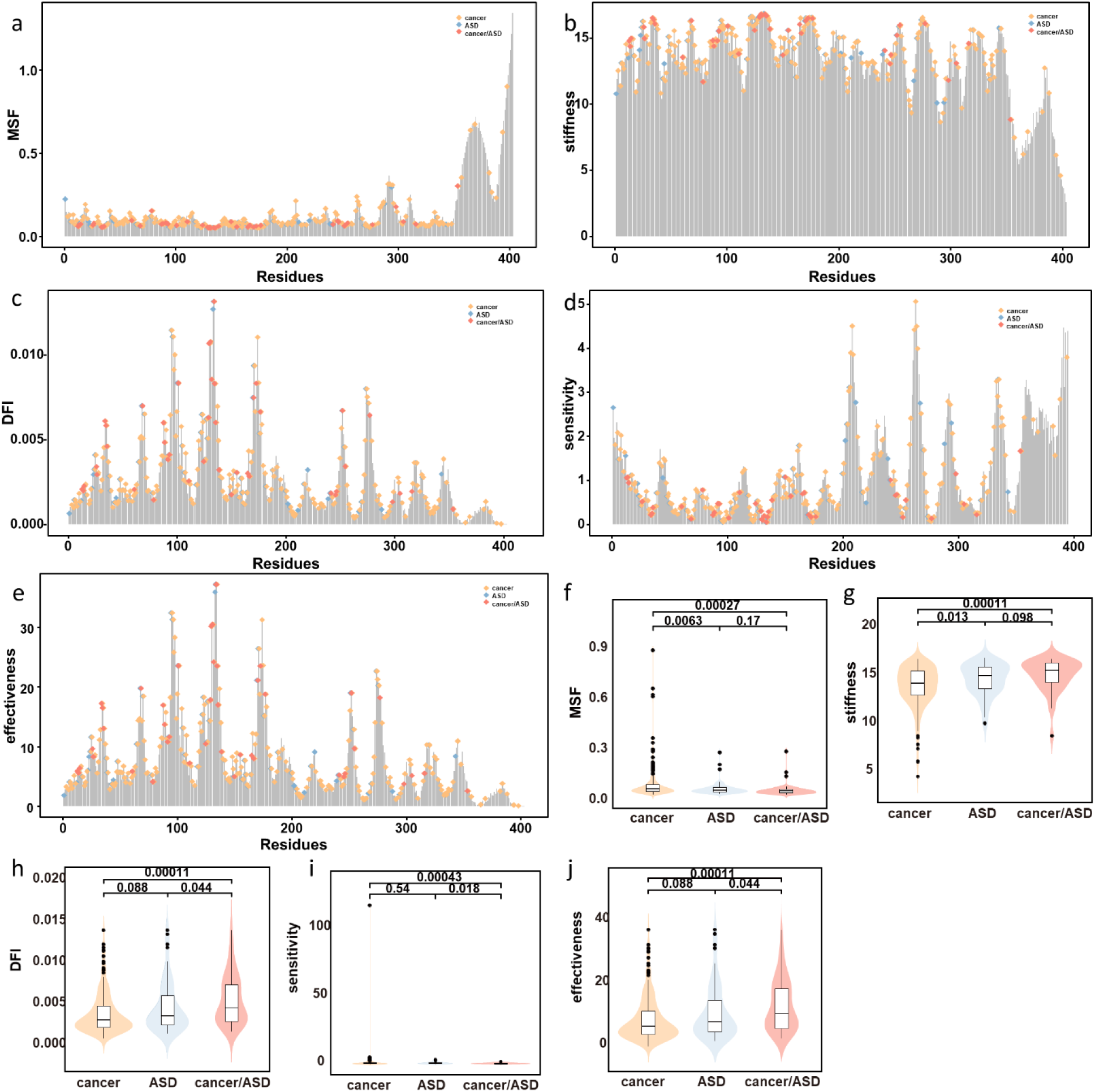
Elastic network model-based features of PTEN mutations. (a) *MSF*, (b) stiffness, (c) *DFI*, (d) sensitivity, (e) effectiveness values, respectively, across PTEN residues (b) Statistical comparison of (f) *MSF*, (g) stiffness, (h) *DFI*, (i) sensitivity, (j) effectiveness, respectively, values among PTEN-cancer, PTEN-ASD, and PTEN-cancer/ASD groups.

Statistical analysis revealed significant differences in *MSF* only between the PTEN-cancer and PTEN-ASD groups (P-value=0.0063) and between PTEN-cancer and PTEN-cancer/ASD groups (P-value=0.00027) (Figure 7f). Similarly, stiffness values showed significant differences only between the PTEN-cancer and PTEN-ASD groups (P-value=0.013), and between PTEN-cancer and PTEN-cancer/ASD groups (P-value=0.00011) (Figures 7g). The *DFI* analysis demonstrated significant differences across all three groups (Figures 7h), highlighting unique flexibility profiles associated with PTEN-cancer/ASD mutations. Sensitivity values differed significantly between the PTEN-ASD and PTEN-cancer/ASD groups (P-value=0.018) and between PTEN-cancer and PTEN-cancer/ASD groups (P-value=0.00043), with no significant difference observed between PTEN-cancer and PTEN-ASD groups (Figures 7i). Effectiveness values also exhibited significant differences across all three groups (Figures 7j).

These findings suggest that PTEN-cancer/ASD mutations confer distinct dynamic properties to the protein, particularly in regions critical for its catalytic and regulatory functions. The observed increases effectiveness, *DFI*, and stiffness may indicate altered allosteric communication and reduced flexibility, potentially impacting the interactions of PTEN with substrates and regulatory proteins. Conversely, the decreases in *MSF* and sensitivity reflect a more rigid structural environment, potentially compromising PTEN’s functional dynamics. Overall, the unique ENM-derived features associated with PTEN-cancer/ASD mutations highlight their substantial impact on PTEN’s structural and functional integrity, providing insight into the molecular mechanisms underlying their dual roles in cancer and NDDs.

### Machine Learning Classification and integrated score for PTEN-cancer/ASD mutations

To effectively distinguish PTEN-cancer/ASD mutations from PTEN-cancer and PTEN-ASD mutations, ROC analysis was conducted on 15 molecular features. The contribution of each feature to phenotype differentiation was evaluated. Based on these selected features, classification models were developed to differentiate among the PTEN-cancer, PTEN-ASD, and PTEN-cancer/ASD groups. Comparison of the AUC values for the 15 features reveled that Δ*CC* demonstrated the highest discriminatory power across the three groups (Figures 8a-c, Figure S3). Specifically, for distinguishing between the PTEN-cancer and PTEN-ASD groups,Δ*CC* achieved an AUC of 0.9311, indicating high reliability. However, when distinguishing PTEN-cancer/ASD from PTEN-cancer groups and PTEN-ASD groups, the AUC values for Δ*CC* were only 0.7747 and 0.6883, respectively, highlighting challenges in achieving precise differentiation.

**Figure 8.**
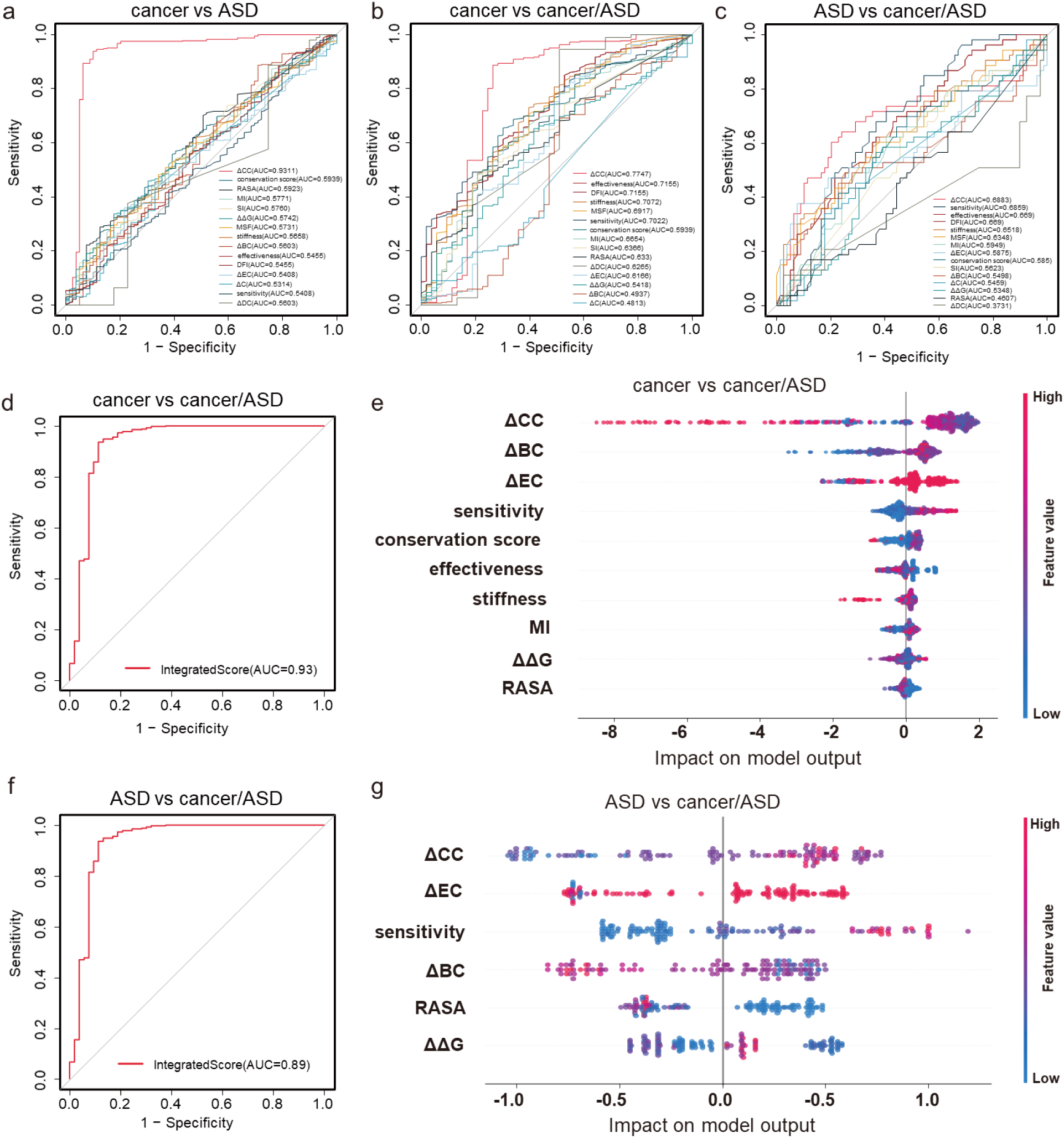
Feature selection to distinguish different phenotypes. The ROC curve and model interpretation plot for single feature selection for distinguishing the PTEN-cancer group from the (a) PTEN-ASD group and (b) PTEN-cancer/ASD group. (c) The ROC curve and model interpretation plot for single feature selection for distinguishing PTEN-ASD group from PTEN-cancer/ASD group. (d) The ROC curve and (e) model interpretation plot for the ML model used to distinguish the PTEN-cancer group from the PTEN-cancer/ASD group. This model was generated by the optimal feature combination composed of Δ*CC*, Δ*BC*, Δ*EC*, sensitivity, conservation score, effectiveness, stiffness, *MI*, *RASA*, and *ΔΔG*. (f) The ROC curve and (g) model interpretation plot for the ML model used to distinguish the PTEN-ASD group from the PTEN-cancer/ASD group. This model included multiple features consisting of Δ*CC*, Δ*EC*, sensitivity, Δ*BC*, *RASA*, and *ΔΔG*.

Machine learning methods were employed to iteratively select multiple features until the AUC exceeded 0.8, demonstrating the model’s ability to effectively distinguish the PTEN-cancer/ASD group from the other two groups (Figures 8d and 8f). The final feature set revealed that distinguishing PTEN-cancer from PTEN-cancer/ASD mutations could be achieved using a combination of Δ*CC*, Δ*BC*, Δ*EC*, sensitivity, conservation score, effectiveness, stiffness, *MI*, *RASA*, and *ΔΔG* (Figure 8e). And, distinguishing PTEN-ASD from PTEN-cancer/ASD mutations could be accomplished using a combination of Δ*CC*, Δ*EC*, sensitivity, Δ*BC*, *RASA*, and *ΔΔG* (Figure 8g). Notably, the selected feature sets consistently included at least one feature from sequence-based, structural, and dynamical properties. These highlights that the model comprehensively accounts for all potential contributors to the distinct allosteric properties associated with these different phenotypes. The top four features critical for distinguishing the PTEN-cancer/ASD group were Δ*CC*, Δ*BC*, Δ*EC*, and sensitivity, which are crucial for revealing the allosteric properties of the PTEN-cancer/ASD mutations.

Additionally, the developed ML model can generate an IS for multi-type variants, facilitating a more granular classification of similar phenotypes. The IS confirmed that PTEN-cancer/ASD mutations, including R130Q, R173C^13, 14, 23^, Y155C^13, 43^, and C136R^13, 23^, all exhibited scores greater than 0.5, validating their classification as PTEN-cancer/ASD mutations and further supporting the model’s effectiveness and accuracy.

Previous studies have suggested that PTEN-ASD mutations predominantly affect local regions, especially within the PD, causing local allosteric effects that disrupt communication and functionality within the domain^13^. In contrast, PTEN-cancer mutations typically induce global allosteric effects, influencing the conformation and function of the entire protein via long-range structural communication pathways. However, the specific nature of communication (local, global, or both) for PTEN-cancer/ASD mutations remains unclear. Additionally, we also identified several PTEN-cancer/ASD mutations with exhibited high IS compared to PTEN-cancer and PTEN-ASD mutations (Figure S4, Table S7), including G132A, R130P, T131I and Y68N. These mutations are believed to exhibit significant allosteric properties. To investigate further, MD simulations were performed on these four PTEN-cancer/ASD mutations to explore their dynamic effects in greater detail.

### Dynamics of PTEN-cancer/ASD mutations: local open conformation coupled with long-range allosteric communications

Through MD simulations (see Text S2), we observed that the four PTEN-cancer/ASD mutations — Y68N, R130P, T131I, and G132A (Figures S5 and S6) — differentially affected the flexibility of both local and distal amino acid residues within the PTEN protein. All four mutations altered the flexibility of key functional loops (WPD, P, and TI loops), transitioning them from a closed to an open conformation. Specifically, the R130P, T131I, and G132A mutations in the P loop notably increased the distance between residues R130 and D92 (Figure 9a). This disruption of the compact WT conformation was especially evident in the TI loop, which exhibited increased flexibility and a marked deviation from its native closed conformation, suggesting structural destabilization that could affect distal regions (Figure S7). MD simulations were conducted for 500 *ns* for each of the four mutations and the WT PTEN, eventually reaching relative stability (Figure 9b). The Y68N mutation not only affected Region 1 and Region 2 in the PD but also influenced the flexibility of residue A79, transmitting the allosteric signal to a more distal region within the same domain. This change was reflected in the reduced stability of residue A79 and its surrounding area, indicating a structural alteration in the distal region of the PD (Figures 9c and 9d). Similarly, the R130P mutation significantly impacted the stability of residues H93 and T167, located on the WPD loop and TI loop, respectively. The mutation at position 130 residue within the P loop, further confirmed the functional synergy between the WPD loop, TI loop, and P loop (Figures 9e and 9f). The T131I mutation notably affected the stability of Region0 (residue: L42-V45) centered around residue E43 in the same domain (Figures 9g and 9h). In contrast, the G132A mutation transmitted the signal to the distal residue G143, leading to changes in the stability of the surrounding region (Figures 9i and 9j).

**Figure 9.**
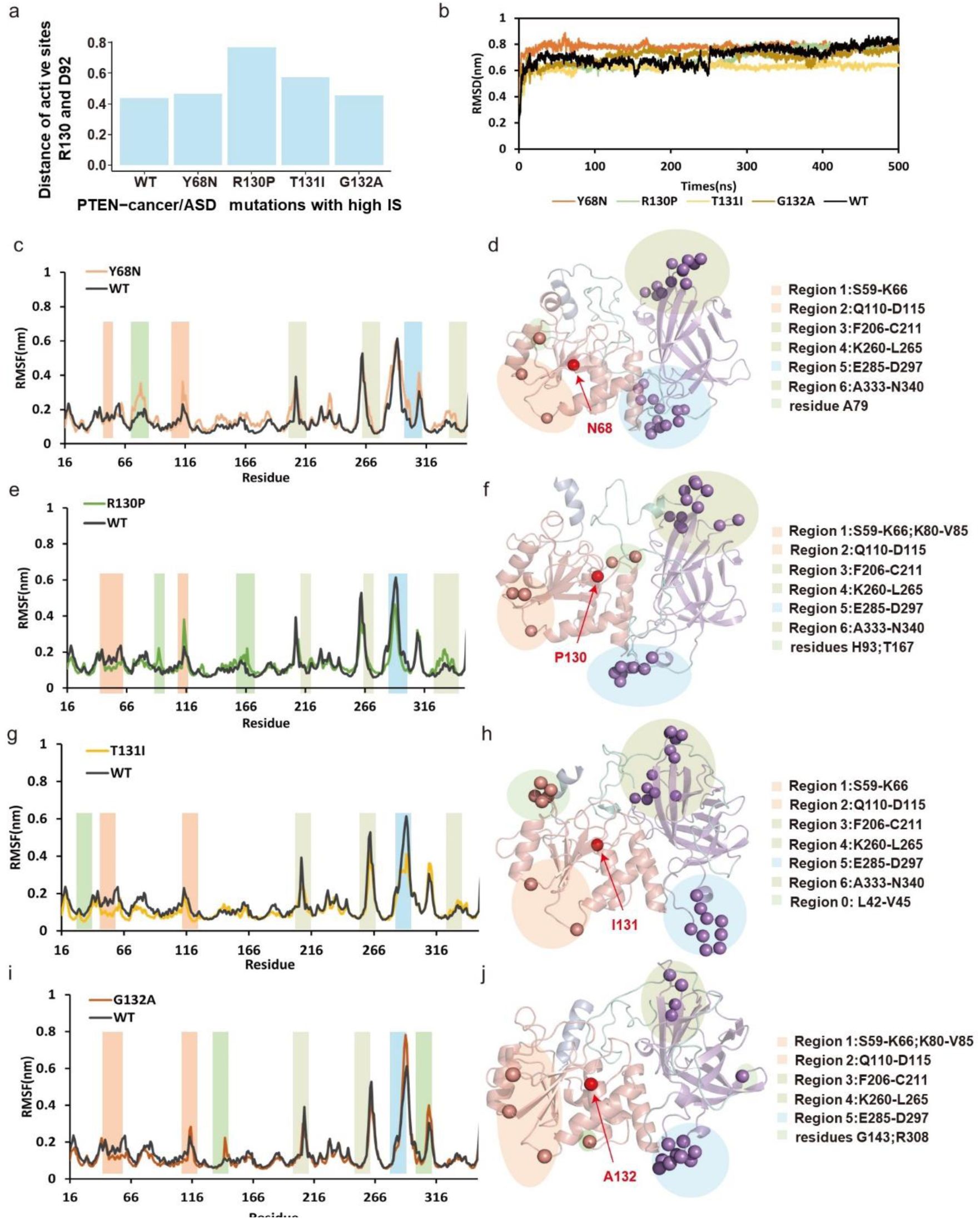
Conformational effects in WT PTEN system and high-scoring PTEN-cancer/ASD systems. (a) Distance of active sites D92 and R130 with PTEN-cancer/ASD mutations. (b) RMSF results for residues in the WT PTEN and mutants Y68N, R130P, T131I, and G132A. RMSF for (c) Y68N, (e) R130P, (g)T131I and (i) G132A and Spatial distribution Spatial distribution of the residues with significant flexibility changes due to the (d) Y68N, (f) R130P, (h)T131I and (j) G132A mutations. Detailed information about these regions is provided in the Figure S1.

These results suggest that PTEN-cancer/ASD mutations with high scores in the P loop region not only affect the stability of the active site but also influence the stability of key functional loop regions. Furthermore, these mutations influence the stability of residues surrounding the mutation site and potentially transmit their effects to more distal regions, suggesting that such high-scoring mutations exhibit stronger allosteric effects. The residues with the highest flexibility changes in the Y68N, R130P, T131I, and G132A mutants were L295, T286, T286, and K289, respectively, indicating potential allosteric communication between the mutation sites and these regions. Utilizing a dynamic network model-based shortest path algorithm, our results demonstrate that mutants exhibit shorter allosteric pathways, further supporting our hypothesis that PTEN-cancer/ASD mutations induce a clear allosteric tendency (Figure 10).

**Figure 10.**
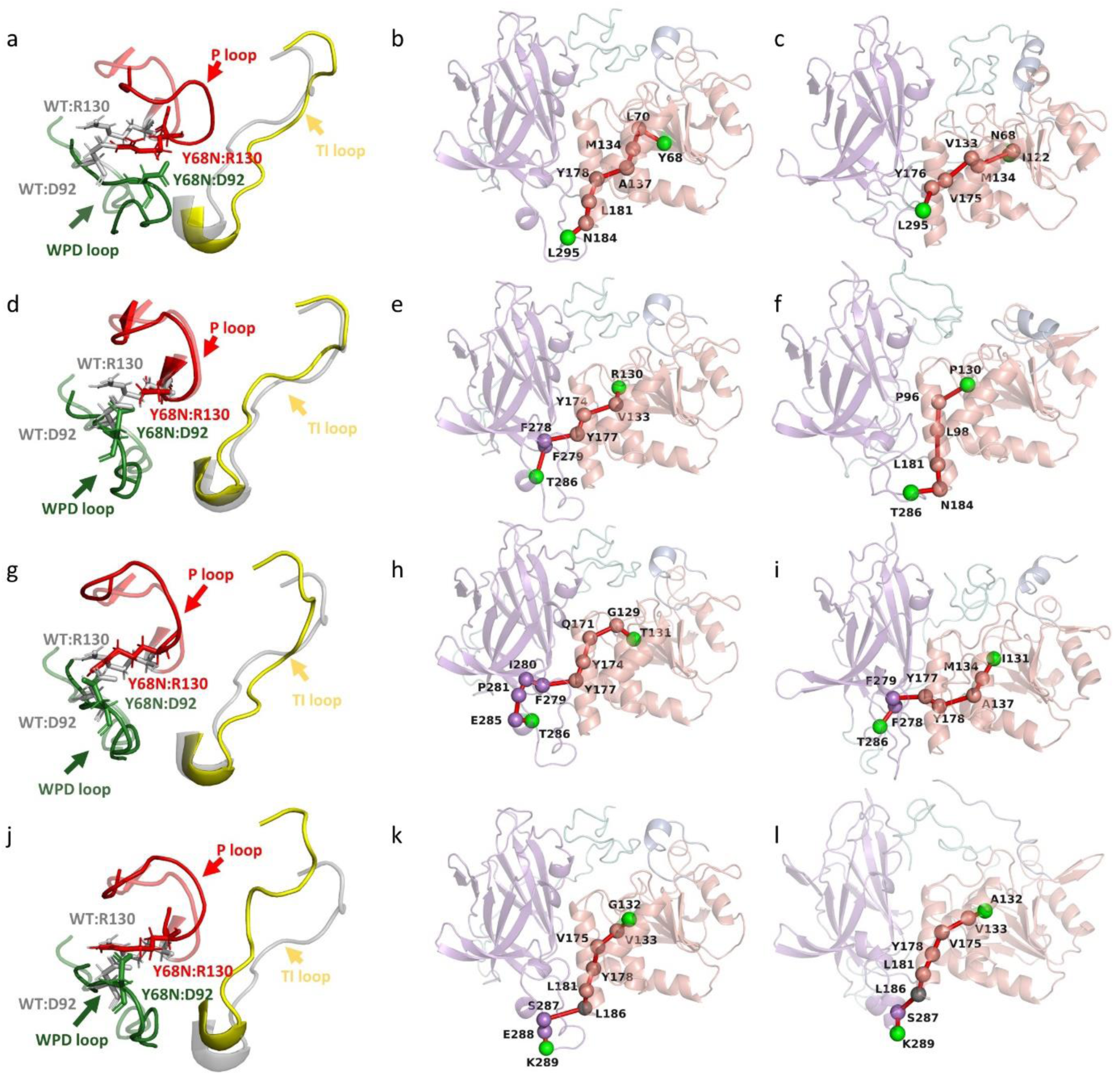
Conformational pathways of high-IS PTEN-cancer/ASD mutations selected by the model. (a) Local conformational changes in the active site of WT-Y68 and mutant Y68N. Conformational pathway of (b) WT-Y68, (c) mutant Y68N, with the starting point at residue 68 and the endpoint at residue L295. (d) Local conformational changes in the active site of WT-R130 and mutant R130P. Conformational pathway of (e) WT-R130, (f) mutant R130P, with the starting point at residue 130 and the endpoint at residue T286. (g) Local conformational changes in the active site of WT-T131 and mutant T131I. Conformational pathway of (h) WT-T131, (i) mutant T131I, with the starting point at residue 131 and the endpoint at residue T286. (j) Local conformational changes in the active site of WT-G132 and mutant G132A. Conformational pathway of (k) WT-G132, (l) mutant G132A, with the starting point at residue G132 and the endpoint at residue K289.

In the WT Y68, the shortest path from residue Y68 to residue L295 required passing through the L181 and N184 regions to transmit the signal to the distal region. However, following the Y68N mutation, the active site adopts an open conformation (Figure 10a), and the shortest path bypasses the L181 and N184 regions, instead choosing a shorter, more direct route to transmit the signal to the distal region (Figures 10b and 10c). This alteration is likely due to the opening of the active site region. Furthermore, after the Y68N mutation, the shortest path no longer traverses residue L70, but shifts to a path passes through the P loop. Similarly, the R130P mutation also resulted a shorter, more direct pathways following the opening of the region (Figure 10d). Before the R130 mutation, the optimal shortest path predominantly passed through Motif 1, reaching the C2D to transmit the allosteric signal to residue T286. After the R130P mutation, the shortest path navigated through the WPD loop and interdomain residues L181 and N184 to transmitting the signal to T286. While the path before and after the mutation was similar, the post-mutation path short, involving fewer residues passing through the WPD loop. This alteration is likely related to the open states of the WPD loop, P loop, and TI loop (Figures 10e and 10f). The T131I and G132A mutants also exhibited shortened paths, with signals passing through residues Y177 and L186 (Figures 10g-l). These findings suggest that PTEN-cancer/ASD mutations with high IS not only perturb the stability of the active site but also propagate this disturbance to distal regions, inducing conformational changes in those areas.

In these pathways, we observed that, in the wild-type PTEN, signal propagation largely depends on interdomain interfaces and key surrounding residues such as L181, N183, and N184. However, PTEN-cancer/ASD mutations tend bypass these interdomain connection points, transmitting the signal to distal regions. Molecular feature analysis further reveals that residues V133, M134, Y177, and Y178 exhibit relatively high effectiveness and low sensitivity, serving as key sites for structural signal propagation between the mutated domain and interdomain regions.

## 4. Conclusions

Predicting the specific phenotype of missense mutations is critical for advancing precision medicine. PTEN, a well-studied protein, is associated with diverse mutations linked to multiple phenotypes, including cancer and ASD. While numerous studies have focused on distinguishing PTEN-cancer and PTEN-ASD mutations, PTEN-cancer/ASD mutations present distinct molecular characteristics, integrating both oncogenic and neurodevelopmental effects within a shared structural framework. This dual functionality complicates the understanding of their specific pathways and raises two key questions: (1) Why do certain PTEN mutations promote both cancerous and neurodevelopmental phenotypes in different individuals or contexts? (2) How do these dual-faceted mutations structurally modulate PTEN’s activity, resulting in variability in disease outcomes?

In this study, we developed an integrative ML model combined with structural dynamics and network-based analyses to systematically evaluate PTEN-cancer/ASD mutations. Previous ML approaches have focused on classifying PTEN mutations based on their molecular consequences on protein structure and function, distinguishing PTEN-cancer, ASD, and non-pathogenic mutations ^17^. Our model builds on this foundation by incorporating four network and dynamic-based features^47^, introduce four network/dynamics-based features— Δ*BC*, Δ*CC*, Δ*EC*, and sensitivity—which complement the conventional set of molecular descriptors (e.g., *ΔΔG*). Compared to the commonly used import features, Δ*CC* and *ΔΔG*, these new features collectively form an “intrinsically accessible spectrum of modes of motions” that enables adaptation to adapt to different environments and interactions while maintaining its structural fold and functional integrity. Among these dynamics features, allosteric signaling effectiveness emerged as a key factor. Overall, our model underscores the critical role of structural dynamics in predicting the phenotypic effects of missense mutations. This approach led to the development of a novel integrative score, which not only classifies PTEN mutations but also ranks the likelihood of mutations belonging to the PTEN-cancer/ASD group.

To enhance the interpretability of ML predictions and the underlying molecular mechanism, we performed MD simulations combined with network analysis on selected PTEN-cancer/ASD mutations. Several comprehensive computational studies have previously leveraged MD simulations for analyze PTEN mutations. For example, previous investigations have examined the conformational alterations of the catalytic core in the PD^24^, while other studies have demonstrated the allosteric effects of PTEN missense mutations^48^. By integrating network analysis, MD simulations have revealed distinct differences in allosteric regulation between PTEN-ASD and PTEN-cancer mutations^21^. These differences arise from the coupled interplay of CTT phosphorylation dynamics. Notably, two key studies employing MD-based computational approaches provided valuable insights into the genotype-phenotype relationships of PTEN-cancer and PTEN-ASD mutations^13, 14^. Their findings reveal that 1) PTEN-cancer mutations induce long-range communication pathways that span the inter-domain interface, while maintaining a closed conformation at the active site, 2) PTEN-ASD mutations cause localized destabilization restricted to the PD, leading to partial opening of the active site. Building on these studies, we applied a similar MD-network-based method to investigate the dynamic effects of PTEN-cancer/ASD mutations. Our results demonstrate that PTEN-cancer/ASD mutations exhibit coupled dynamics characterized by local conformational changes and long-range allosteric communications. Specifically, these mutations induce a partially open conformation at the P loop, altering PTEN’s structural dynamics. Accordingly, we hope that our work will expand the genotype-phenotype map of PTEN mutations, shedding light on the unique pathways associated with PTEN-cancer/ASD mutations.

To enhance the of our ML models, the structural dynamics analysis provided valuable insights into the molecular mechanisms underlying PTEN-cancer/ASD mutations. Among the most significant features, Δ*CC* and Δ*BC* capture signatures of local conformational changes, while effectiveness and sensitivity elucidate how mutations act as molecular drives of allosteric communication pathways. Furthermore, our results highlight the potential of ML models in precision medicine, particularly in the context of PTEN-cancer/ASD mutations. One promising strategy involves targeting mutations located at active sites or orthosteric sites by designing effective modulators to control local conformational change. Another viable approach is the development of allosteric drugs that modulate the long-range dynamics caused by these mutations. Interestingly, mutations in other drug targets^49, 50^ show similar dynamics profiles, with both localized effects and long-range allosteric changes. This observation suggests the combining orthosteric and allosteric drugs could form a transferable therapeutic paradigm for treating multi-phenotypic mutations^51^.

## ASSOCIATED CONTENT

### Data Availability Statement

The codes for ML model and the data for MD simulations are available at https://zenodo.org/records/14636196.

## Supporting Information

Text S1. Detailed information of structural modeling, molecular feature calculations, and MD simulations.

Text S2. The results of the MD simulations.

Figure S1. PTEN structure and mutation mapping.

Figure S2. Sequence conservation scoring results of PTEN.

Figure S3. Feature correlation analysis.

Figure S4. Distribution of IS for PTEN-cancer/ASD mutations.

Figure S5. MD trajectory convergence and reproducibility.

Figure S6. RMSF comparison among independent MD replicates

Figure S7. Conformational effects in wild-type PTEN and high-IS PTEN-cancer/ASD mutants.

Table S1. Overview of collected PTEN missense mutation data. These data were curated from public databases, ensuring a broad and representative set of pathogenic variants for subsequent analyses.

Table S2. RASA values computed by PASIS for PTEN. These data were used to evaluate whether mutations were buried, partially exposed, or exposed, providing important structural context for PTEN-cancer, PTEN-ASD, and PTEN-cancer/ASD mutations.

Table S3. PTEN sequence-based features computation results. These data were used to examine how sequence conservation and co-evolutionary differ across the three PTEN mutation groups.

Table S4. Folding free energy (ΔΔG) computation results. These data reveal the thermodynamic impacts of mutations and highlight notable hotspots in PTEN-cancer/ASD variants.

Table S5. PTEN amino acid contact energy network features. These network-based features are pivotal in distinguishing local vs. global structural perturbations among PTEN variants.

Table S6. Elastic network model-based dynamic features. These data elucidate the dynamic consequences of PTEN-cancer, PTEN-ASD, and PTEN-cancer/ASD mutations.

Table S7. Machine learning integrated scores for PTEN-cancer/ASD classification. This set of scores corroborates the ML-based findings discussed in the main text and reveals high-IS mutations that merit further structural and functional analysis.

## AUTHOR INFORMATION

### Corresponding Authors

Guang Hu - MOE Key Laboratory of Geriatric Diseases and Immunology, Suzhou Key Laboratory of Pathogen Bioscience and Anti-infective Medicine, Department of Bioinformatics and Computational Biology, School of Life Sciences, Suzhou Medical College of Soochow University, Suzhou 215123, China, Fei Xiao - Department of Bioinformatics and Computational Biology, School of Life Sciences, Suzhou Medical College of Soochow University, Suzhou 215123, China

### Authors

Miao Yang - Department of Bioinformatics and Computational Biology, School of Life Sciences, Suzhou Medical College of Soochow University, Suzhou, 215213, China

Jingran Wang - Department of Bioinformatics and Computational Biology, School of Life Sciences, Suzhou Medical College of Soochow University, Suzhou, 215123, China

Ziyun Zhou - Department of Bioinformatics and Computational Biology, School of Life Sciences, Suzhou Medical College of Soochow University, Suzhou, 215123, China

Wentian Li - Department of Bioinformatics and Computational Biology, School of Life Sciences, Suzhou Medical College of Soochow University, Suzhou, 215123, China

Gennady Verkhivker - Department of Biomedical and Pharmaceutical Sciences, Chapman University School of Pharmacy, Irvine 92618, California, United States

### Author Contributions

^#^M. Y. and J. W. contributed equally. The manuscript was written through contributions of all authors. All authors have given approval to the final version of the manuscript.

### Notes

The authors declare no competing financial interest.

## ACKNOWLEDGMENT

This work was supported by the National Natural Science Foundation of China (32271292), the Project of MOE Key Laboratory of Geriatric Diseases and Immunology (JYN202404), and a Project Funded by the Priority Academic Program Development (PAPD) of Jiangsu Higher Education Institutions. This research was also funded by the National Institutes of Health under Award 1R01AI181600-01 and Subaward 6069-SC24-11 to Gennady Verkhivker.

