## Supplementary material for "Decoding Mechanisms of PTEN Missense Mutations in Cancer and Autism Spectrum Disorder using Interpretable Machine Learning Approaches": Text S1

**MATERIALS AND METHODS**

**Structural modeling.** The full-length PTEN protein structure was obtained from the AlphaFold Protein Structure Database^1, 2^. The quality and confidence of the predicted structure were assessed using the AlphaFold per-residue confidence score ($\text{pLDDT}$) Additionally, structural alignment between the AlphaFold-predicted PTEN structure and the experimentally resolved structure (PDB ID: 1D5R) from the Protein Data Bank^3^ was performed using TM-align^4^, a structure alignment toll integrated within the I-TASSER suite^5^.

**Molecular features computation**

**Relative Accessible Surface Area (**$\boldsymbol{RASA}$**).** The relative Accessible Surface Area ($RASA$) is a normalized measure of the solvent accessibility of an amino acid residue within a protein^6^. The $RASA$ of each PTEN residue was analyzed using PSAIA^7^ and classified into three categories based on their $RASA$ values: buried ($RASA \leq5\%$), partially exposed ($5\% < RASA \leq30\%$), and exposed ($RASA > 30\%$).

$$RASA=ASA_{residue}/ASA_{max}$$

(1)

where $ASA_{residue}$ is the accessible surface area ($ASA$) of the amino acid residue in protein and $ASA_{max}$ is the $ASA$ of the residue in an extended state. The calculation of $RASA$ is bases on DSSP^3^.

**Conservation and coevolution-based features.** Residue conservation within the PTEN protein was assessed using the ConSurf server^8^, which provides scores ranging from 1 to 9, with higher values indicating greater conservation. Furthermore, a refined multiple sequence alignment (MSA) of the PTEN protein was retrieved from the ConSurf server for Shannon information entropy $S(i)$and mutual information$(MI)$ calculations. In this study, we characterized both variable and conserved regions of PTEN, estimating the conservation levels for each amino acid site, thereby highlighting regions that are critical for its biological activity. Conservation levels were estimated for each amino acid site, highlighting regions critical for PTEN’s biological activity. $S(i)$ is calculated using the formula:

$$S\left( i \right)=-\sum_{ai=1}^{20} P\left( ai \right)\log P\left( ai \right)$$

(2)

where $P(xi,yj)$ represents the relative frequency of the $i^{th}$ amino acid at position $i$. The $S(i)$ typically ranges from 0 to 3 where 0 indicates no variability and higher values signify greater uncertainty. The co-evolution score measures the degree of interdependence between residues during evolution, a common method for assessing co-evolution is based on mutual information $(MI)$, which quantifies the statistical dependency between residues and is calculated as:

$$I\left( i,j \right)=\sum_{xi=1}^{21} \sum_{jy=1}^{21} P\left( xi,yj \right)\log\frac{P\left( xi,yj \right)}{P\left( xi \right)P\left( yj \right)}$$

(3)

where $P(xi,yj)$ is the joint probability of observing amino acid types $x$ and $y$ at sequence positions $i$ and $j$, respectively, while is the marginal probability of type $x$ at position $i$. The value of $I(i,j)$ varies within the range$[0,Imax]$, reflecting the degree of correlation between residue pairs^9^. By analyzing the average $MI$ for each residue, we can assess the co-evolution of mutations. The calculation of $S(i)$and $MI$ was performed by Evol^10^.

**Gibbs free energy.** Structural stability changes induced by mutations were investigated using FoldX^11^. Each PTEN missense variant and its corresponding mutant structures were automatically generated to calculate the difference in folding Gibbs free energy $(\Delta\Delta G$) for each PTEN mutant. The positive and negative $\Delta\Delta G$ values indicate a decrease and increase in protein stability, respectively, while higher absolute values of $\Delta\Delta G$ signify a larger effect on protein stability^12^.

**Network-bases features.** We utilized the NACEN R package^13^ to calculate the amino acid network parameters derived from the constructed Amino Acid Contact Energy Networks (AACENs)^14^. In AACEN, proteins are transformed into graph structures, with each amino acid residue as a node and whether there are edges between the nodes depends on the environment-dependent residue contact energy (ERCE) between the residues. The ERCE is calculated as follows:

$${ERCE}_{vu}=-ln\frac{N_{vu}N_{00}C_{v0}C_{u0}}{N_{v0}N_{u0}C_{vu}C_{00}}$$

(4)

where$N_{vu}$represents the number of contacts between amino acid residues $i$ and $j$ observed in the actual protein structure. $N_{00}$ denotes the number of contacts between two amino acids in a randomized state. $N_{v0}$ and $N_{u0}$ are the total numbers of exposures of amino acids $i$ and $j$ in a particular environment and $C_{v0},C_{u0},C_{vu},C_{00}$ account for the local environmental context in which the amino acids are situated, ensuring that environmental influences on contact probabilities are appropriately adjusted. Additionally, pattern recognition on geometric features was conducted to build unweighted networks for PTEN-cancer, PTEN-ASD, and PTEN-cancer/ASD mutations. Network topology was analyzed using centrality parameters^15^. Degree Centrality ($DC$) is a local parameter that counts the number of edges connecting a node to its neighboring nodes. Nodes with high degree centrality are often viewed as hubs within the network, and it is defined as:

$$DC\left( v \right)=\frac{k_{v}}{n-1}$$

(5)

where $k_{v}$ is the number of connections for node $v$, and $n$ is the total number of nodes in the network. The Clustering Coefficient ($C$) measures the density of connections between a node and its neighboring nodes. It is calculated using the connectivity relationships in the network, providing insight into the local density of connections. Nodes with a high clustering coefficient indicate a relatively high number of connections among their neighbors, forming more triangular structures, which are more compact and stable compared to quadrilateral structures. It is calculated by:

$$C\left( v \right)=\frac{2T\left( v \right)}{deg\left( v \right)\left( \deg\left( v \right)-1 \right)}$$

(6)

where $T(v)$ represents the total number of connection pairs among neighbors, and $deg(v)$ indicates the number of neighbors of node $v$. Betweenness Centrality ($BC$) is a global parameter related to the shortest paths between different nodes in a network. Nodes with high betweenness centrality indicate their significant role in network communication and may serve as critical bottlenecks in signal transmission. It can be calculated as follows:

$$BC\left( v \right)=\sum_{s\neq v\neq t} \frac{\sigma_{st}\left( v \right)}{\sigma_{st}}$$

(7)

where $\sigma_{st}$is the total number of shortest paths from node $s$ to node $t$, and$\sigma_{st}\left( v \right)$is the number of those paths that pass through node $v$. Closeness Centrality ($CC$) is another global parameter that reflects the shortest paths between nodes in the network. A higher closeness centrality indicates that communication from that node to others is relatively easier. It is calculated by:

$$CC\left( v \right)=\frac{1}{\sum_{u\neq d} d\left( v,u \right)}$$

(8)

where $k_{v}$ is the number of connections for node $v$, and $n$ is the total number of nodes in the network. Eigenvector Centrality ($EC$) measures the quality of a node's connections and its overall importance within the network. This metric helps identify key nodes and potential core components of the network, defined as:

$$EC\left( v \right)=\frac{1}{\lambda}\sum_{j} A_{ij}x_{j}$$

(9)

where $A_{ij}x_{j}$ represents the adjacency matrix of the network, $\lambda$ is the largest eigenvalue, and $x_{j}$ denotes the centrality of node $j$.

**Dynamics network features.** To analyze the dynamic effect of mutations, the Elastic Network Model (ENM)^16, 17^ was employed, comprising the Gaussian Network Model (GNM) and the Anisotropic Network Model (ANM). The total potential energy in the GNM and ANM describes the system's response to deviations from equilibrium. For GNM, the total potential energy ($V_{GNM}$) of a system with N nodes is expressed as:

$$V_{GNM}=\frac{\gamma}{2}\sum_{i=1}^{N} \sum_{j=1}^{N} \Gamma_{ij}\left( \Delta R_{i}\cdot\Delta R_{j} \right)$$

(10)

where $\gamma$ is the uniform spring constant, $\Delta R_{i}\cdot\Delta R_{j}$are the displacements of nodes from their equilibrium positions, and $\Gamma_{ij}$ represents the ${ij}^{th}$ elements of the $N\times N$ Kirchhoff matrix that define connectivity between nodes, calculated as:

$$\Gamma_{ij}=\left\{ \begin{aligned} -1,i\neq j\cap R_{ij}\leq r_{c} \\ 0,i\neq j\cap R_{ij}>r_{c} \\ -\sum_{i,i\neq j} \Gamma_{ij},i=j \end{aligned} \right.$$

(11)

In this study, the cutoff distance $r_{c}$ between protein nodes were 7 $Å$. ANM involves more complex, direction-aware potential energy calculations, enabling the study of anisotropic deformations that more accurately mimic real protein motions. For ANM, the potential energy ($V_{ANM}$) takes into account the directionality of displacements and is given by:

$$V_{ANM}=\frac{\gamma}{2}\sum_{i=1}^{N} \sum_{j=1,j\neq i}^{N} H_{ij}\left( R_{i}-R_{j} \right)^{2}$$

(12)

where $H_{ij}$ denotes the Hessian matrix elements representing the second derivatives of the potential energy with respect to atomic coordinates, capturing the directional interaction between nodes $i$ and $j$. The Hessian matrix $H_{ij}$ is written as:

$$H_{ij}=\left[ \begin{matrix} \frac{\partial^{2}V}{\partial x_{i}{\partial x}_{j}} & \frac{\partial^{2}V}{\partial x_{i}{\partial y}_{j}} & \frac{\partial^{2}V}{\partial x_{i}{\partial z}_{j}} \\ \frac{\partial^{2}V}{\partial y_{i}{\partial x}_{j}} & \frac{\partial^{2}V}{\partial y_{i}{\partial y}_{j}} & \frac{\partial^{2}V}{\partial y_{i}{\partial z}_{j}} \\ \frac{\partial^{2}V}{\partial z_{i}{\partial x}_{j}} & \frac{\partial^{2}V}{\partial z_{i}{\partial y}_{j}} & \frac{\partial^{2}V}{\partial z_{i}{\partial z}_{j}} \end{matrix} \right]$$

(13)

where $V$ is harmonic potential calculated by:

$$V=\frac{\gamma}{2}\left( R_{i}-R_{j} \right)^{2}$$

(14)

where $R_{i}$ and $R_{j}$ are the positions of the nodes. For dynamics features^18^, effectiveness and sensitivity are derived from the ANM framework using the Perturbation Response Scanning (PRS) matrix^19^, measure the protein's efficiency in performing its function and its response to structural changes, respectively. stiffness, calculated based on the ANM, is derived by averaging the columns of the stiffness matrix, reflecting the protein's resistance to structural changes. Dynamic Flexibility Index ($DFI$) and Mean-Square Fluctuations ($MSF$) are computed using the GNM, where $DFI$ measures the flexibility of each node, and $MSF$ quantifies the magnitude of atomic fluctuations around equilibrium positions, offering insights into protein flexibility. Here, the GNM and ANM calculations were performed by ProDy^20^.

**Molecular Dynamics simulations**

**Molecular Dynamics Setup and Parameterization.** All-atom molecular dynamics (MD) simulations were performed with the GROMACS software package (version 5.1.4) ^21^, employing the Amber99SB-ILDN force field^22^. Initial coordinates for wild-type (WT) PTEN were energy-minimized and equilibrated under NVT (100 $ps$) and NPT (100 $ps$) ensembles at 303.15 $\text{K}$, followed by a 500 $ns$ production run. We used a 2 $fs$ integration time step, LINCS constraints on hydrogen-containing bonds, and Particle-Mesh Ewald (PME) for long-range electrostatics ($\text{cutoff = 12 Å}$). Periodic boundary conditions were applied in all directions.

**Mutant Construction and Replicates.** Single-point PTEN-cancer/ASD mutants (Y68N, R130P, T131I, G132A) were generated using FoldX^23^, followed by the same minimization/equilibration procedure. To ensure reproducibility, we conducted three independent replicates (each 500 $ns$) for both WT and mutant systems, yielding a total simulation time of 7.5 $\mu s$. The replicate trajectories were merged for statistical analyses once convergence was confirmed (Figures S5 and S6).Finally, GROMACS, VMD^24^, and PYMOL^25^ were employed for the computational analysis and visualization of the MD trajectories, enabling the investigation of the structural and dynamic properties of both WT and mutants.
