## Supplementary material for "Decoding Mechanisms of PTEN Missense Mutations in Cancer and Autism Spectrum Disorder using Interpretable Machine Learning Approaches": Text S2

**RESULTS**

**Structural modeling.** In this study, we used the predicted structure of PTEN from AlphaFold Protein Structure Database^1, 2^. The confidence of the AlphaFold-predicted structures was assessed using the $\text{pLDDT}$, where both the PD and the C2D showed high confidence levels ($\text{pLDDT}$ > 80), supporting the reliability of these regions. Additionally, the average RMSD on backbone α-carbon (Cα) atoms was used to quantify the similarity between protein tertiary structures, we compared it against the experimental structure available in the PDB (PDB ID: 1D5R). The comparison revealed RMSD of 1.56 Å and a TM-score of 0.79, indicating a substantial degree of similarity and structural reliability between the AlphaFold-predicted structure and the experimentally resolved segments.

**MD simulations.** Through the integrated model screening, mutations with high Integrated Score (IS) were found to cluster around the P loop region of the PD (Figure S7a), consistent with previous studies^3^. To investigate the conformational impact of these high-IS mutations on different regions, four single-point mutant models were constructed: Y68N, R130P, T131I, and G132A. Each mutant system, along with the wild-type structure, was subjected to 500 $ns$ of MD simulations, during which all systems reached a relatively stable state. The flexibility trends of the Cα atoms in each system were subsequently calculated (Figure S7b). Overall, the RMSF profiles for the four mutant systems were generally consistent with the WT, with higher RMSF values predominantly localized in some unstable loop regions.

As shown in Figures S7c-f, all four mutant systems affected two regions within the phosphatase domain: **Region 1** (S59–K66) and **Region 2** (Q110–D115). These two regions lie in close spatial proximity and exhibited marked flexibility changes in both the residues and their surrounding environments. Notably, the stability of the Region 2 decreased, whereas Region 1 became more stable, suggesting a potential synergistic relationship between these two regions. Furthermore, the mutations transmitted their effects to another domain, introducing pronounced flexibility changes in the **Region 3** (F206–C211), **Region 4** (K260–L265), and **Region 5** (E285–Q297). Importantly, all four mutations induced a jump-like change in the flexibility of the Region 3. The perturbation was also transmitted to the **Region 6** (A333–N340), an inter-domain region that is spatially close to Region 3 and Region 4, despite being distant in the primary sequence. This inter-domain region (Region 6) exhibited decreased stability. Of particular interest, Region 4 corresponds to the CBR3 loop (previously reported in the literature^3-6^) and is functionally associated with membrane binding. Perturbations within this loop can induce conformational changes at the active site, potentially leading to an isomeric activation mechanism in PTEN. Taken together, these observations highlight the capacity of PTEN-cancer/ASD mutations with high IS to propagate local perturbations into more distal regions, thereby exerting broader allosteric effects on the protein’s structural and functional integrity.
