## Supplementary material for "Decoding Mechanisms of PTEN Missense Mutations in Cancer and Autism Spectrum Disorder using Interpretable Machine Learning Approaches": Figure S

**Supporting Figures**

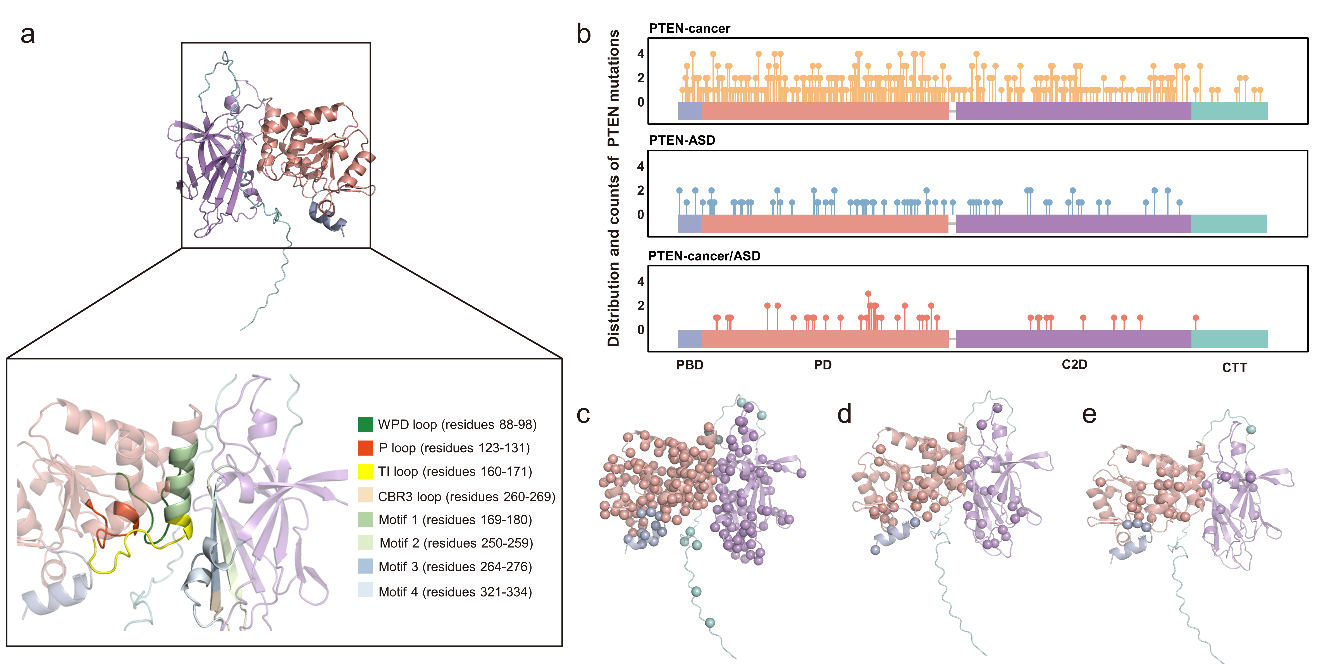

**Figure S1. PTEN structure and mutation mapping.** The full-length AlphaFold-predicted structure of PTEN shows an N-terminal domain (PBD), residues 1-15 (blue); a catalytic phosphatase structural domain (PD), residues 16-185 (red), and interaction interfaces. (C2D), residues 190-35 (purple); a C-terminal tail (CTT), residues 351-403 (green), and a membrane-bound structural domain. Detailed annotation of functional motifs and loops: WPD loop (residues 88–98), P loop (residues 123–131), TI loop (residues 160–171), CBR3 loop (residues 260–269), and four structural motifs (residues 169–180, 250–259, 264–276, and 321–334). (b) Distribution of PTEN-associated mutations across different domains. (c) PTEN-cancer mutations, (d) PTEN-ASD mutations and (e) PTEN-cancer/ASD mutations map on the three-dimensional structure.

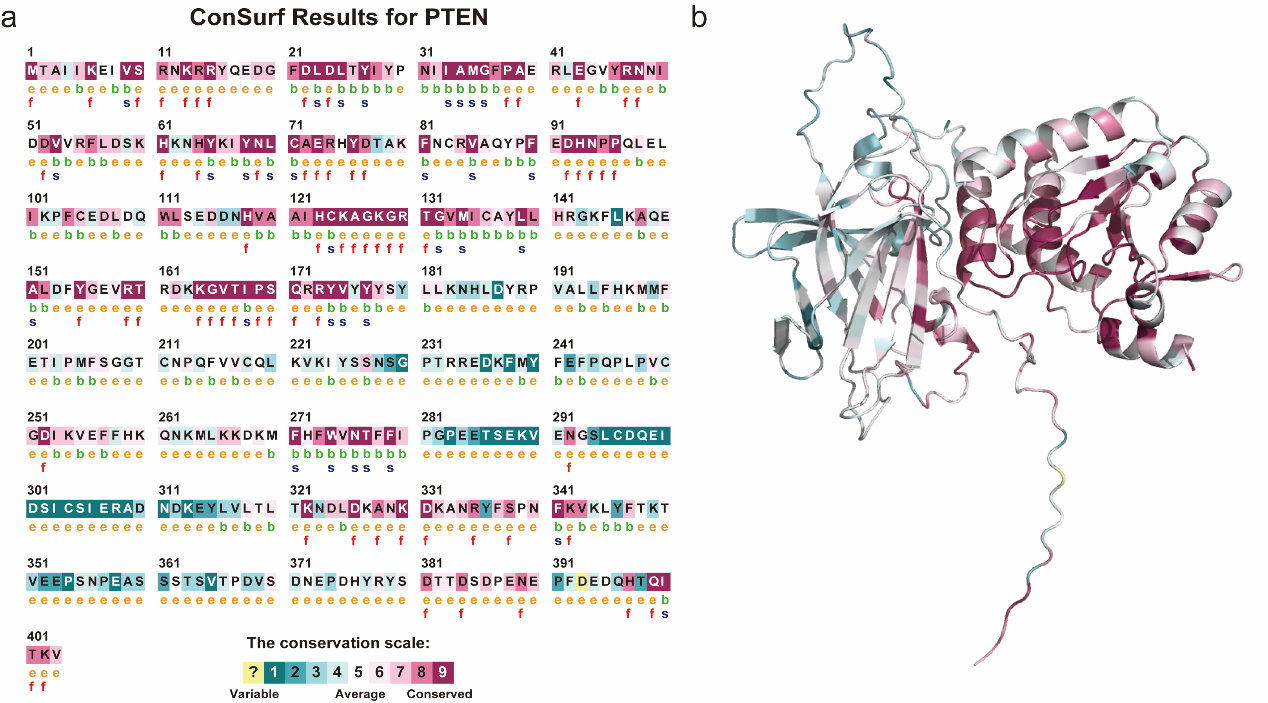

**Figure S2. Sequence conservation scoring results of PTEN on the Consurf server.** The conservation scale from Variable (score 0, yellow) to Consolidated (score 9, red). (a) PTEN sequence with conservation score. The meaning of the letters below the sequence is: e-An exposed residue according to the neural network algorithm. b - A buried residue according to the neural network algorithm. f - A predicted functional residue (highly conserved and exposed). s - A predicted structural residue (highly conserved and buried). X - Insufficient data - the calculation for this site was performed on less than 10% of the sequences. Disease-causing mutations are usually located in conserved positions in proteins. (b) Conservation of three-dimensional structural sequences. The sites with high conservation scores are distributed in the phosphatase domain.

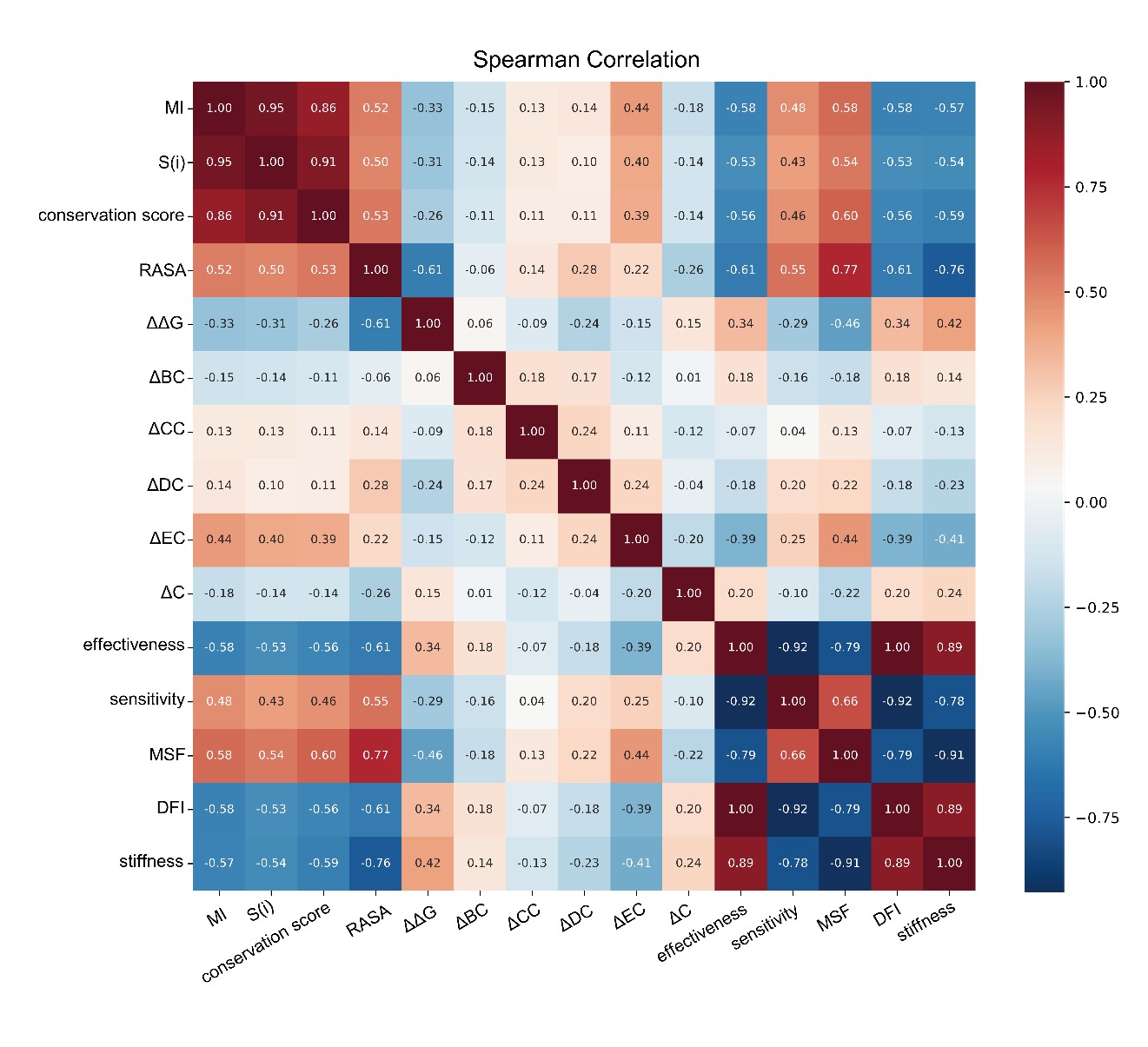

**Figure S3. Feature correlation analysis.** The gradient from blue to red indicates a shift in correlation coefficients among the features, ranging from negative (blue) to positive (red). Different shades represent varying correlation strengths: deeper blue signifies a stronger negative correlation, while deeper red indicates a stronger positive correlation.

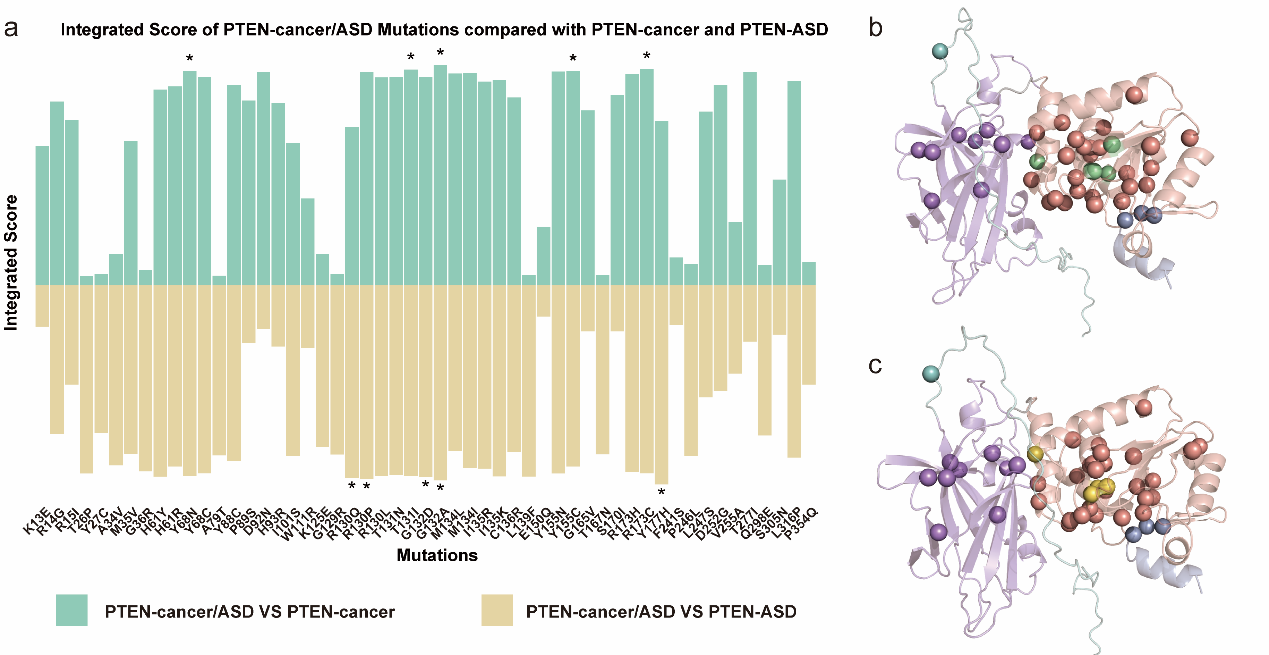

**Figure S4.** **Distribution of IS for PTEN-cancer/ASD mutations.** (a) Comparison of the IS between PTEN-cancer mutations (green), PTEN-ASD mutations (yellow), and PTEN-cancer/ASD mutations (red). Asterisks indicate the top five mutations with the highest IS. (b) Mapping of PTEN-cancer/ASD mutations onto the 3D structure, overlaid with PTEN-cancer mutations (green). (c) Mapping of PTEN-cancer/ASD mutations onto the 3D structure, overlaid with PTEN-ASD mutations (yellow). We observed that PTEN-cancer/ASD mutations with high IS, as predicted by the machine learning model, predominantly cluster in the PD, consistent with our previous analyses. Notably, several PTEN-cancer/ASD mutations exhibited higher IS values than those observed in PTEN-cancer or PTEN-ASD groups, including: G132A (PTEN-cancer/ASD vs. PTEN-cancer: 0.9437390; PTEN-cancer/ASD vs. PTEN-ASD: 0.8377885), R130P (PTEN-cancer/ASD vs. PTEN-cancer: 0.9145012; PTEN-cancer/ASD vs. PTEN-ASD: 0.8309291), T131I (PTEN-cancer/ASD vs. PTEN-cancer: 0.9249051; PTEN-cancer/ASD vs. PTEN-ASD: 0.8195578), and Y68N (PTEN-cancer/ASD vs. PTEN-cancer: 0.9190731; PTEN-cancer/ASD vs. PTEN-ASD: 0.8191917).

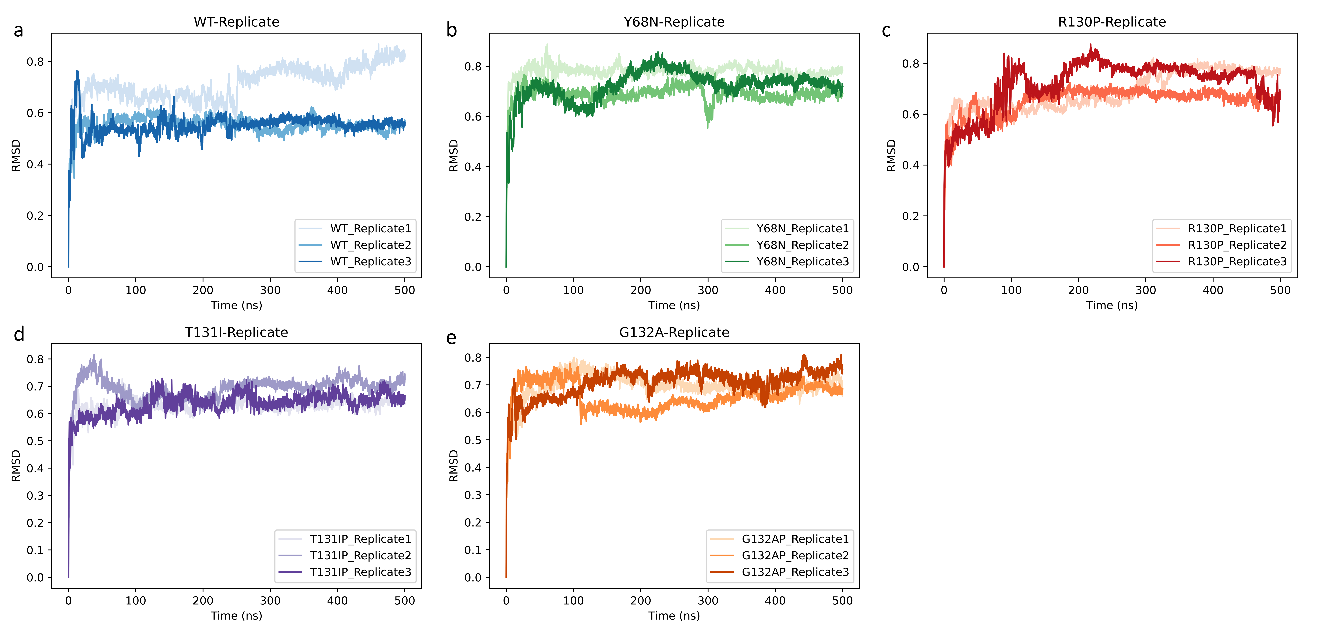

**Figure S5.** **MD trajectory convergence and reproducibility**. (a) RMSD vs. time for WT PTEN. Three independent 500 $ns$ replicates (Replicate 1,2,3) are shown in different colors, illustrating the convergence behavior and overall stability of the WT system during the production runs. (b–e) RMSD vs. Time for Each PTEN-cancer/ASD Mutant (Y68N, R130P, T131I, and G132A). Similar to the WT, each mutant system was simulated in triplicate, demonstrating that all replicates reach stable RMSD plateaus within the first ~100–200 $ns$. In all cases, the RMSD traces plateau at comparable values, indicating that each system converges to a stable conformational ensemble. The overlap among the three replicates for each system underscores the reproducibility of the simulation setup and parameter choices.

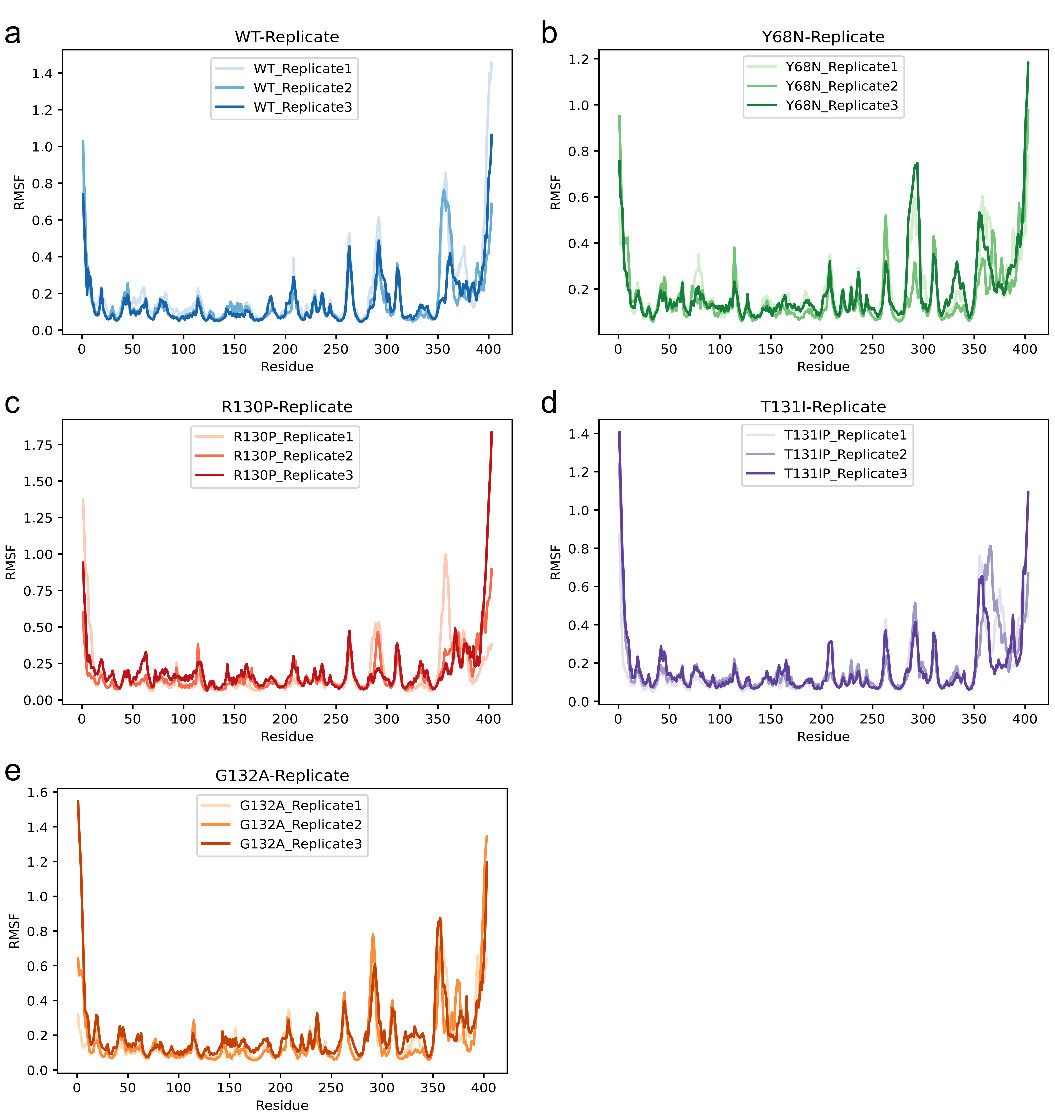

**Figure S6****.** **RMSF comparison among independent MD replicates.** (a) RMSF profiles of wild-type PTEN. Averaged over the final 100 $ns$ of the simulation for each of the three replicates, highlighting residue-level flexibility. Shaded error bands (or separate curves) indicate the variation among replicates. (b–e) RMSF Profiles for PTEN-cancer/ASD mutants (Y68N, R130P, T131I, and G132A). Each panel compares the three replicates’ RMSF values for a given mutant, illustrating that the major flexibility peaks (e.g., loop regions) are consistently observed. Overall, the similarity in RMSF trends among the three replicates supports the notion that each mutant system reaches a convergent sampling. Regions displaying higher RMSF values generally correspond to loops or surface-exposed segments, whereas the protein core remains comparatively rigid.

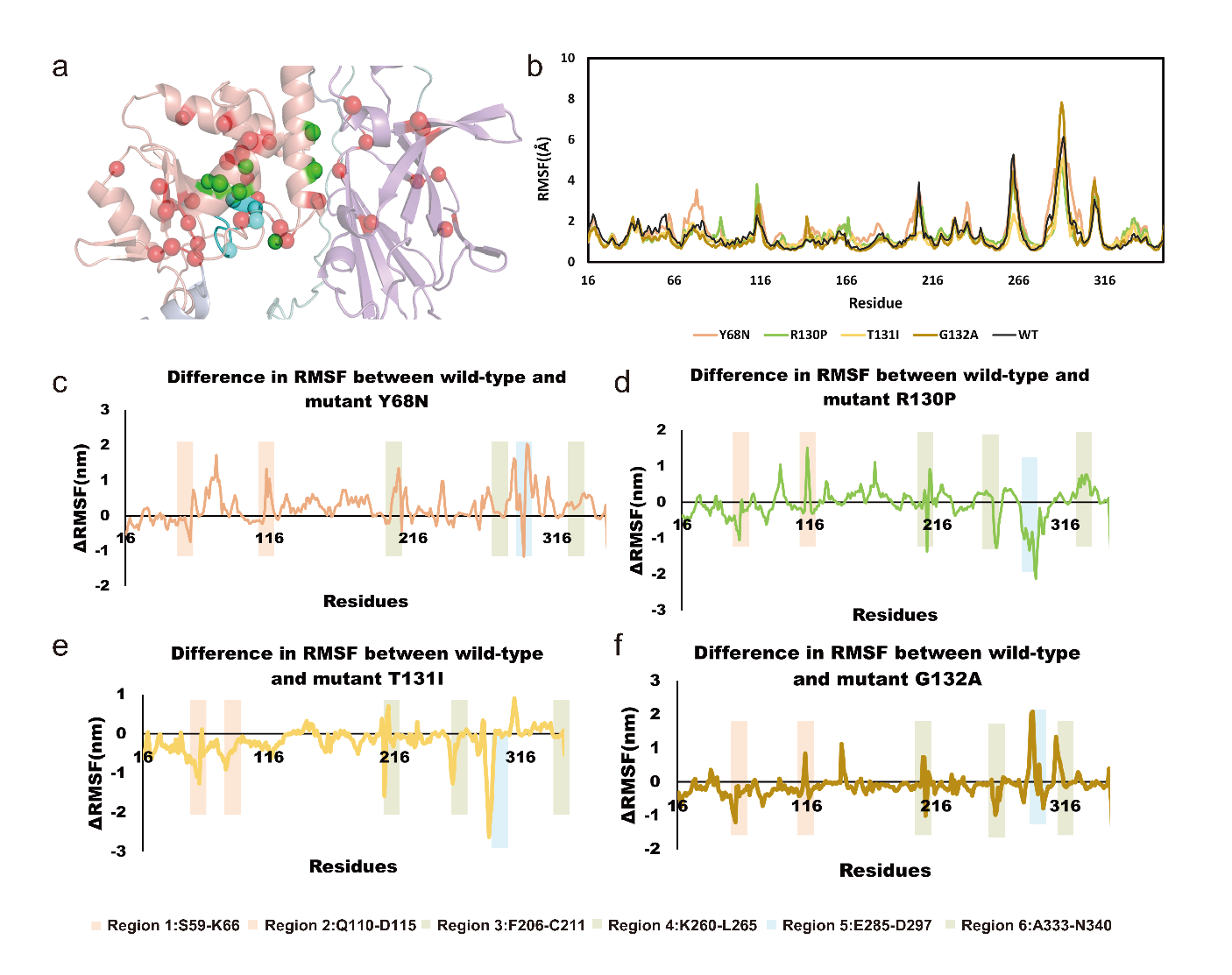

**Figure S7.** **Conformational effects in wild-type PTEN and high-IS PTEN-cancer/ASD mutants.** (a) Distribution of PTEN-cancer/ASD mutations, with the P loop highlighted in blue. PTEN-cancer/ASD mutations are shown in red, and high- IS PTEN-cancer/ASD mutations identified by the integrated model are shown in green. (b) RMSF profiles of WT PTEN and four mutant systems (Y68N, R130P, T131I, G132A). RMSF values are plotted against residue numbers to facilitate comparison of residue-level flexibility. Differences in RMSF (ΔRMSF) between wild-type PTEN and specific mutants: (c) Y68N, (d) R130P, (e) T131I, and (f) G132A. Positive and negative ΔRMSF values indicate increased or decreased flexibility, respectively. Highlighted shaded regions (Region 1: S59–K66, Region 2: Q110–D115, Region 3: F206–C211, Region 4: K260–L265, Region 5: E285–D297, Region 6: A333–N340) correspond to structurally or functionally important segments of PTEN.
